# Population genomics of Vibrionaceae isolated from an endangered oasis reveals local adaptation after an environmental perturbation

**DOI:** 10.1101/743146

**Authors:** Mirna Vazquez-Rosas-Landa, Gabriel Yaxal Ponce-Soto, Jonás A. Aguirre-Liguori, Shalabh Thakur, Enrique Scheinvar, Josué Barrera-Redondo, Enrique Ibarra-Laclette, David S. Guttman, Luis E. Eguiarte, Valeria Souza

## Abstract

**Background:** In bacteria, pan-genomes are the result of the evolutionary “tug of war” between selection and horizontal gene transfer (HGT). High rates of HGT increase the genetic pool and the effective population size, resulting in open pan-genomes. In contrast, selective pressures can lead to local adaptation by purging the variation introduced by HGT, resulting in closed pan-genomes and clonal lineages. In this study, we explored both hypotheses elucidating the pan-genome of Vibrionaceae isolates after a perturbation event in the endangered oasis of Cuatro Ciénegas Basin (CCB), Mexico, and looking for signals of adaptation to the environments in their genomes.

**Results:** We obtained 42 genomes of Vibrionaceae distributed in six lineages, two of them did not showed any close reference strain in databases. Five of the lineages showed closed pan-genomes and were associated to either water or sediment environment; their high effective population size (*N_e_*) estimates suggest that these lineages are not from a recent origin. The only clade with an open pan-genome was found in both environments and was formed by ten genetic groups with low *N_e_*, suggesting a recent origin. The recombination and mutation estimators (r/m) ranged from 0.0052 to 2.7249, which are similar to oceanic Vibrionaceae estimations; however, we identified 367 gene families with signals of positive selection, most of them found in the core genome; suggesting that despite recombination, natural selection moves the Vibrionaceae CCB lineages to local adaptation purging the genomes and keeping closed pan-genome patterns. Moreover, we identify 598 SNPs associated with an unstructured environment; some of the genes under this SNPs were related to sodium transport.

**Conclusions:** Different lines of evidence suggest that the sampled Vibrionaceae, are part of the rare biosphere usually living under famine conditions. Two of these lineages were reported by the first time. Most Vibrionaceae lineages of CCB are adapted to their microhabitats rather than to the sampled environments. This pattern of adaptation agrees with the association of closed pan-genomes and local adaptation.

## Background

Comparative genomics analyses have shown a wide range of genomic variation within bacteria from different phylogenetic groups [1–3]. This variation range has been explained in part by the wide ecological niche occupied by different bacterial groups [4–8]. Bacterial genomes, in contrast to eukaryotic genomes, usually maintain constant genome sizes [9, 10], suggesting that while horizontal gene transfer (HGT) increases the genome size by adding new genes, selection maintains the genome size by removing deleterious, non-functional or non-useful genes [11–13]. Therefore, bacteria can present very different genomic compositions even within a species, with HGT creating a flexible genome and natural selection purging or maintaining it [10, 14].

Thus, the type of pan-genome is an indication of the evolutionary “tug of war” between selection and HGT. As a prediction, if there are high rates of HGT, the total genetic pool will increase, as well as the effective population size, generating an open pan-genome maintained by natural selection [15]. However, if there is a selective pressure towards local adaptation, the genetic diversity introduced by HGT will be purged, resulting in a closed pan-genome and clonal lineages [14].

To start understanding the reasons why some pan-genomes are open while others are closed, we can analyze the rate and type of recombination. On the one hand, homologous recombination homogenizes populations, keeping them genetically cohesive in a closed pan-genome [16, 17]. On the other hand, non-homologous recombination brings new genetic material, offering new evolutionary opportunities for diversification and generating an open pan-genome [18–21]. Recombination decreases linkage disequilibrium among genes, allowing selection and the related Hill-Robertson effect to operate in specific genes and avoiding the purged of genetic diversity along with the genome [22, 23]. As a result of this diversity, species with higher recombination levels maintain a large historical effective population size [15, 24, 25]. In contrast, highly clonal populations with low or no HGT evolve mostly by mutation and genetic drift, because the efficiency of selection is hampered by the Hill-Robertson effect that also reduces the standing levels of variation in the population and the historical effective population sizes [23, 26].

In this study, we explored the probable role of different evolutionary forces shaping the genetic diversity of Vibrionaceae in the oasis of the Cuatro Ciénegas Basin (CCB), Mexico. CCB is composed of several aquatic systems that have a significant unbalance of the nutrient stoichiometry [27]. Population genetic studies of *Pseudomonas* spp., *Exiguobacterium* spp. and *Bacillus* spp. isolated from CCB ponds and rivers in general have shown low recombination levels [28–30]. These patterns suggest that nutrient constraints in CCB may work as an ecological filter, reducing recombination maybe due to the cost of replicating new DNA, and leading to local adaptation [27, 31, 32].

We tested whether the environmental nutrient constraint would affect the genetic structure of *Vibrio* spp. lineages at CCB. Members of *Vibrio* spp. has been characterized in general as a highly recombinant [33, 34]. We analyzed the genetic structure of Vibrionaceae in a particular site of CCB, Pozas Rojas (Figure 1). This site was the most stoichiometrically unbalanced (N:P 156:1) in our first sampling on 2008 [35]. Later, Pozas Rojas was naturally perturbed with intense rains associated with hurricane Alex in 2010. The runoff detritus and water, caused the nutrients ratios to change from extremely unbalanced stoichiometry to a ratio similar the standard values in the sea (N:P 20:1; compared to the Redfield standard N:P 16:1 values of the sea [36]). Given the change in stoichiometry ratios, we asked the following questions: 1) How did a naturally recombinant lineage like some members of Vibrionaceae respond to this perturbation? 2) Did Vibrionaceae lineages maintained their local adaptation to this unique site by restricting recombination, and maintaining their pan-genomes closed? Alternatively, 3) Is it possible that *Vibrio* spp. developed open pan-genomes with large effective population sizes, similar to the lineages in the ocean to deal with this stoichiometric change? [33, 34].

**Figure 1.**
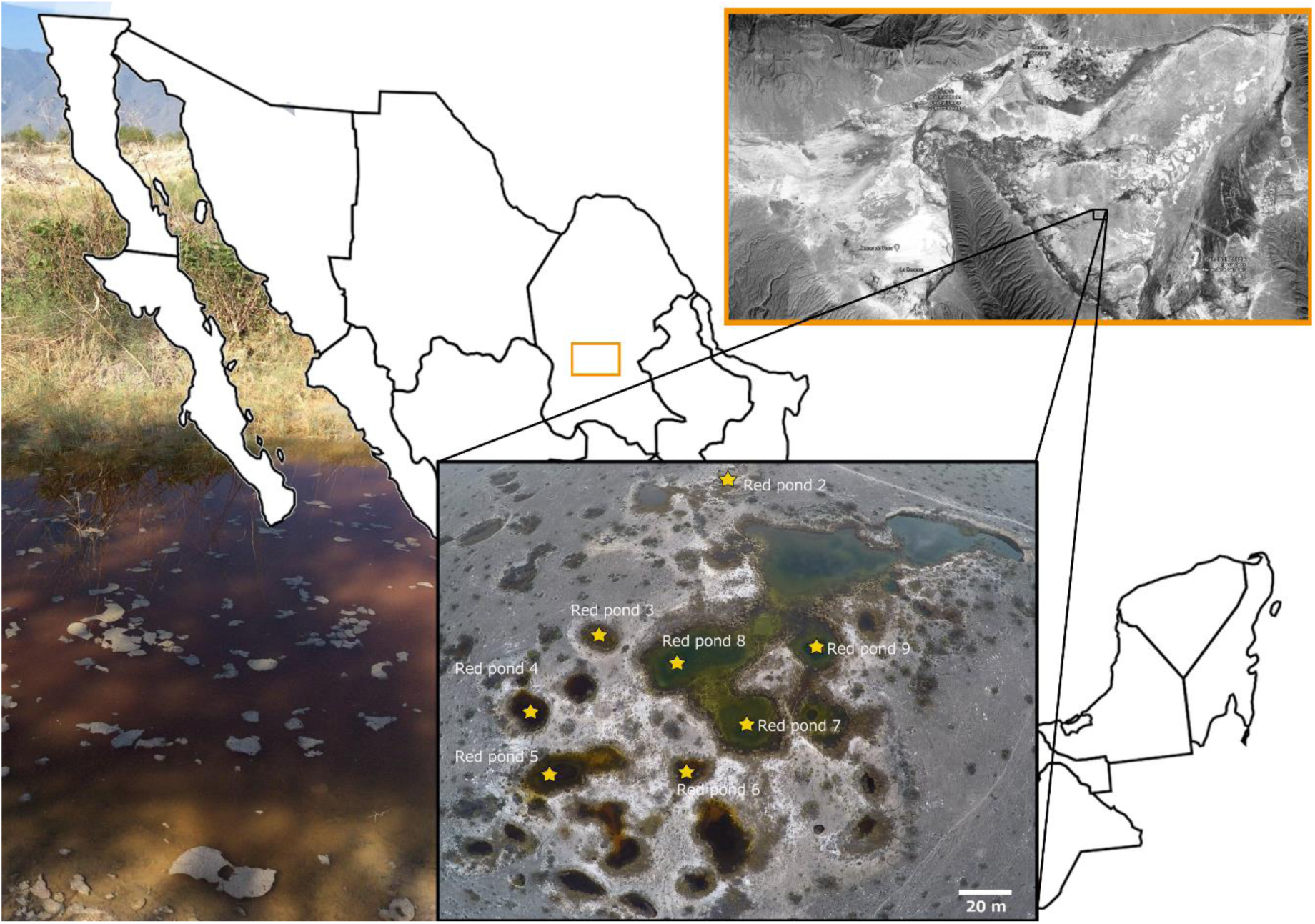
Study site, Pozas Rojas in Los Hundidos within Cuatro Ciénegas Basin, Mexico. Sampling sites are signaled in yellow. Cuatro Ciénegas location is also shown in a map (Pozas Rojas photos were provided by David Jaramillo, a map showing the location of Cuatro Ciénegas Valley was obtained from Google Earth, earth.google.com/web/).

Herein we analyzed the role of the evolutionary forces that have shaped Vibrionaceae at CCB by performing a comparative genomics analysis of five reference and 42 strains isolated from two different local environments (i.e., water and sediments) in perturbed Pozas Rojas. Contrary to what we expected, our results show that most CCB Vibrionaceae lineages had similar levels of recombination compared to their oceanic relatives, and much higher levels of recombination than other genera in the CCB [28–30]. However, since most of the analyzed lineages had closed pan-genomes, we suggest that most of such recombination is homologous. This type of recombination should promote reproductive isolation and generate local adaptation. We did not observe a clear pattern of adaptation to either water or sediment environments, suggesting that there may be other environmental variables that we were not able to measure that could be driving local adaptation among these lineages.

## Results

### Nutrients concentrations

Based on Kruskal-Wallis statistical test, the total nutrient concentrations (Carbon (C), Nitrogen (N), and Phosphorus (P)) of the Pozas Rojas were not significantly different between sample points,(C: *p*= 0.8815; N: *p*= 0.2256 and P: *p*= 0.9624; Additional file 1: Table 1) however, they were statistically significant between type of environment (water vs. sediment) (C: *p*= 3.486e-4; N: *p*= 0.03798 and P: *p*= 3.461e-4).

**Table 1.** Pan-genome metrics of each Vibrionaceae clades isolated from Poza Rojas, CCB.

| Group Clade | Number of CCB genomes included in each clade | Pan-genome metrics |  |  |  | Heaps law parameters |  |
| --- | --- | --- | --- | --- | --- | --- | --- |
|  |  | Core | Flexible | Unique | Total number of genes | Intercept value | Alpha |
| Clade I | 3 | 3617 | 346 | 603 | 4566 | 692.8508 | 1.1293 |
| Clade II | 22 | 1746 | 5770 | 1745 | 9261 | 244.2096 | 0.7913 |
| Clade III | 5 | 2672 | 718 | 324 | 3714 | 658.0634 | 1.6625 |
| Clade IV | 5 | 2055 | 1445 | 180 | 3680 | 2726.7580 | 2.0000 |
| Clade V | 4 | 2853 | 1660 | 1332 | 5845 | 1196.2571 | 1.3109 |
| Clade VI | 3 | 2448 | 3476 | 1028 | 4992 | 3295.5770 | 2.0000 |
| Vibrionaceae all Clades | 47 | 1254 | 14072 | 4795 | 20121 | 2263.7472 | 0.6621 |

The first column shows the Clade ID, next is the number of genomes used for the analysis regarding each clade, followed by the general metrics of pan-genome, and last columns show the heaps values obtained. If alpha >1.0 the pan-genome is considered closed if alpha <1.0 it is considered open.

The proportion of C:N:P was on average 350:9:1 for water, and 258:21:1 for sediment (Additional file 1: Table 2). This ratios indicate a stoichiometric “balance” (i.e., similar to Redfield standard ratios) in Pozas Rojas during 2013, due to higher P availability, compared with the extreme stoichiometric imbalance observed in most of CCB sites, and in particular in Pozas Rojas microbial mat during summer 2008 (i.e., 15,820:157:1)[35], previous to the hurricane Alex perturbation.

**Table 2.** Genetic diversity statistics.

| Clade | | Number of individuals | Number of segregating sites | $\pi$ | $\theta_w$ | Tajima's $D$ | P-value of Tajima's $D$ |
| --- | --- | --- | --- | --- | --- | --- | --- |
| Clade I |  | 3 | 100971 | 0.0164894 | 0.0163978 | 0 | 0 |
| Clade II | All individuals | 22 | 103197 | 0.01148342 | 0.01106029 | 0.15738106 | 0.8582025 |
|  | All individuals in the three larger sub-Clades | 14 | 49946 | 0.00916203 | 0.00613614 | 2.23866585 | 0.02142617 |
|  | Sub-clade G | 4 | 13 | 2.54E-06 | 2.77E-06 | -0.84306779 | 0.77323024 |
|  | Sub-clade D | 6 | 42 | 5.47E-06 | 7.19E-06 | -1.52560731 | 0.02458297 |
|  | Sub-clade A | 4 | 82 | 1.61E-05 | 1.75E-05 | -0.83190864 | 0.8020116 |
| Clade III |  | 5 | 40593 | 0.0051088 | 0.0061293 | -1.27467187 | 0.01772241 |
| Clade IV |  | 5 | 209 | 2.86E-05 | 3.46E-05 | -1.31696234 | 0 |
| Clade V |  | 4 | 34843 | 0.00398639 | 0.00434715 | -0.87361739 | 0.56601856 |
| Clade VI |  | 3 | 204388 | 0.04622002 | 0.04621538 | 0 | 0 |

From left to right are displayed the values for segregation sites, nucleotide diversity ( $\pi$ ) Watterson's theta ( $\theta_w$ ), Tajima's $D$ and Tajima's $D$ $p$ -value. The values were estimated for all six Clades and Sub-clades with 3 or more individuals.

### Phylogenetic Diversity and the environmental association

The phylogenetic relationships of 16S rRNA gene (700 bases) of the 174 cultivated isolates from Pozas Rojas, showed that the strain collection was dominated by Vibrionaceae (63%), followed by Aeromonadaceae (14%) and Halomonadaceae (9.7%; Additional file 1: Table 3). Among Vibrionaceae, we identified two different genera; most strains belong to *Vibrio* spp. (93.6%) and far less to the related *Photobacterium* spp. genus (6.3%).

**Table 3.** Recombination vs. mutation estimates.

| Group Clade | Recombination vs mutation estimates |  |
| --- | --- | --- |
|  | rho/theta | *r/m* |
| Clade I | 0.1036 | 2.7249 |
| Clade II | 0.1171 | 0.5299 |
| Clade III | 0.1498 | 1.1163 |
| Clade IV | 0.1437 | 0.9090 |
| Clade V | 0.0278 | 0.2825 |
| Clade VI | 0.0074 | 0.0052 |
| *P. leiognathi* | 0.0064 | 0.2261 |
| *V. anguillarum* | 0.2889 | 4.0014 |
| *V. ordalii* | 0.0667 | 0.5659 |
| *V. parahaemolyticus* | 0.0025 | 0.1246 |

First column shows the names of the CCB Clades and reference strains used for the calculus. Second and third columns shows the Rho/theta and *r/m* estimates [43].

The aligned sequences were used to construct a maximum likelihood tree with PhyML (Additional file 1: Figure 1). Based on the previous taxonomic assignment, the sequences of *V. alginolyticus, V. parahaemolyticus, V. anguillarum, V. metschnikovii*, and *Photobacterium* spp. were included as references. This analysis reveals seven different cultivated Vibrionaceae lineages in Pozas Rojas.

In order to characterize the relationship between water/sediment environments and Vibrionaceae lineages, we performed an AdaptML analysis [37]. The analysis showed that strains are structured according to the environment where they were isolated, i.e., water or sediment, and not by pond (Additional file 1: Figure 2). While most clades were specialist either to water (higher nutrient condition) or to sediment (lower nutrient condition), the most abundant lineage had no preference for any environment. Based on the AdaptML analysis, we selected 42 isolates for further sequencing; these isolates were chosen as representatives from the different lineages and environments.

**Figure 2.**
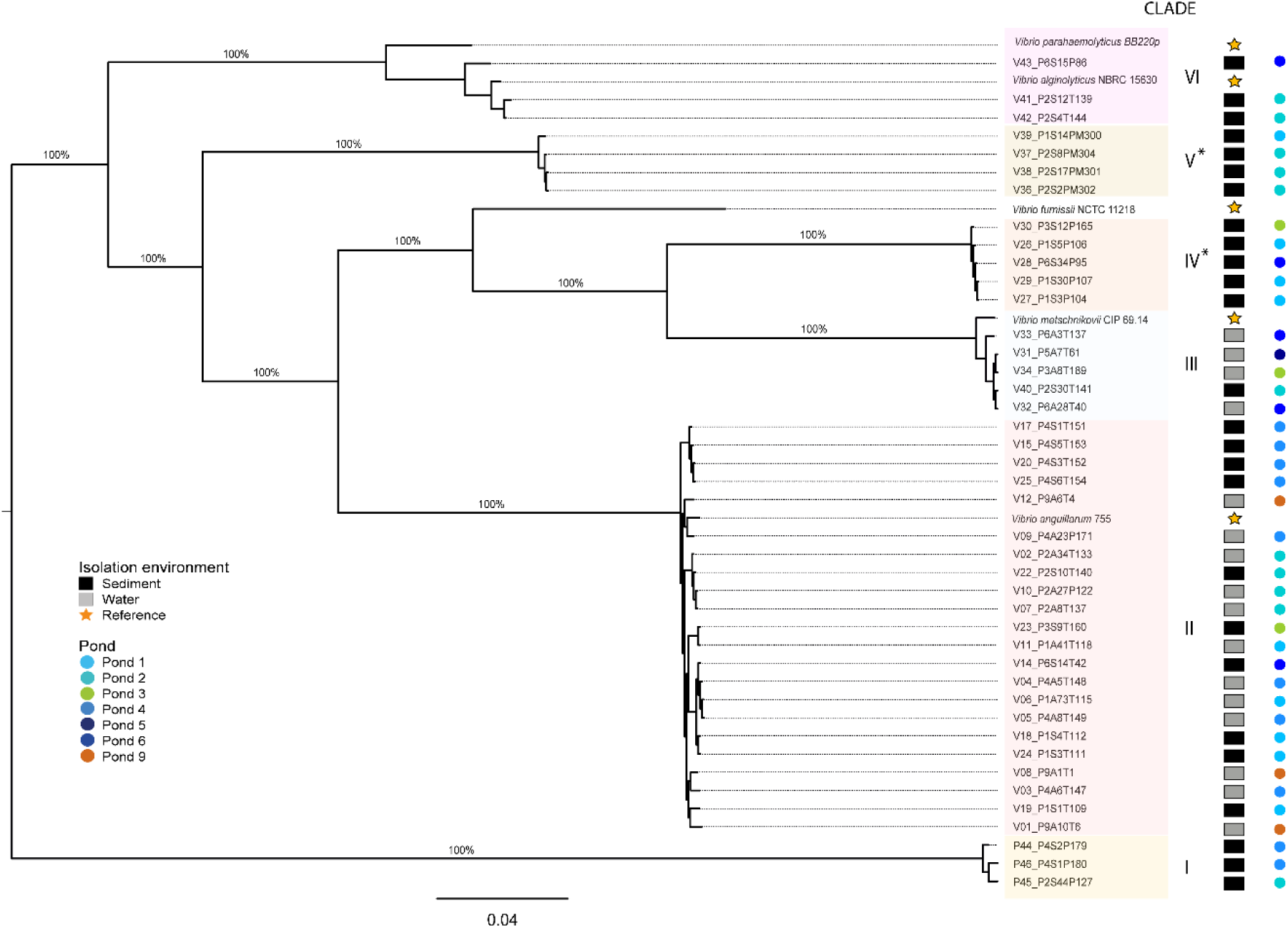
Core gene phylogeny of the 1,254 orthologs. Maximum-likelihood phylogenetic reconstruction of core genes, supporting branch values are shown. Each square represents the isolation environment, water or sediment, while yellow stars indicate reference strains. Circles indicate isolation pond. Clades are distinguished with colors. Clades IV and V which are likely to be exclusive to CCB are highlight with an asterisk.

### Genome features

Among the 39 CCB sequenced *Vibrio* spp. genomes, we found variation in terms of genome size, ranging from 3.1 Mbp to 5.1 Mbp, while the three CCB *Photobacterium* spp. genomes had an average genome size of 4.5 Mbp. Despite this variation, when we compared the CCB strains genomes to their closest reference strain, we found similar genome sizes (Additional file 1: Table 4). Moreover, for each of the assembled genomes, we evaluated their completeness with BUSCO [38]. We found that 92.8% of the genomes contained more than 95% of the 452 near-universal single-copy orthologs evaluated by the program (Additional file 1: Table 5), suggesting that the observed variation in genome sizes could be due to intrinsic characteristics of each strain and not to a sequencing bias.

**Table 4.**
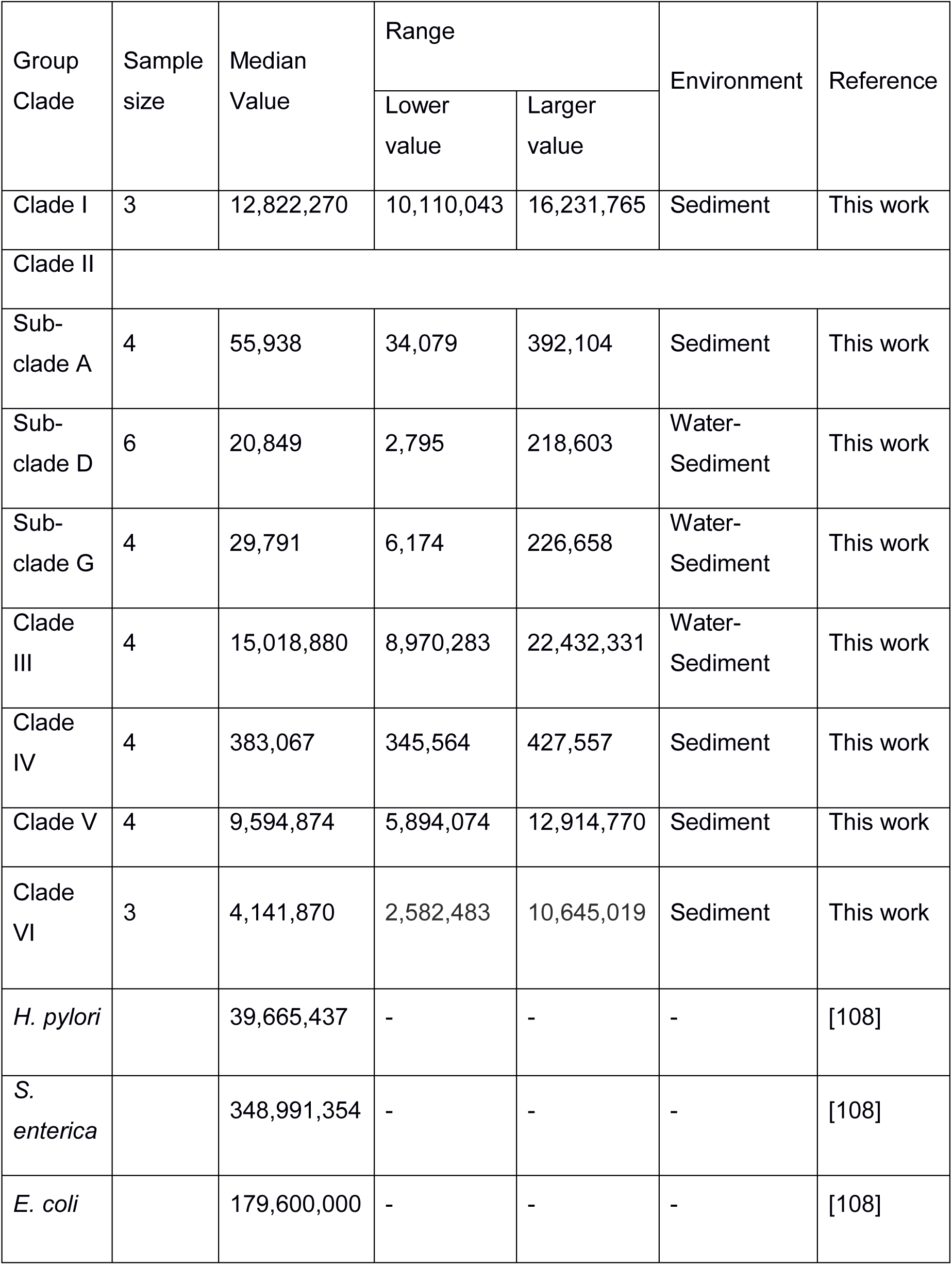
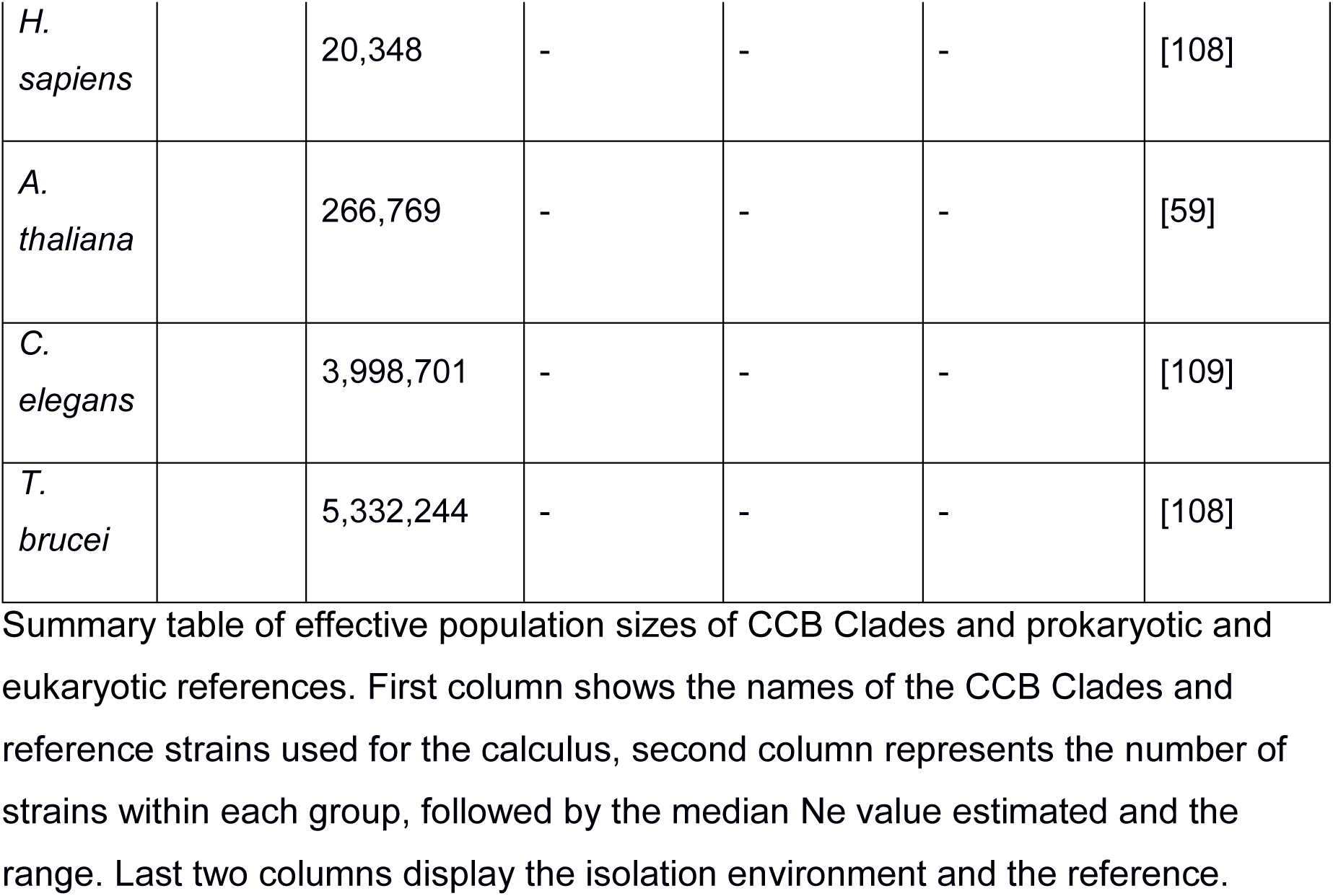
Estimates of effective population sizes obtained through simulations with Fastsimcoal2 [44, 96].

| Group<br>Clade | Sample<br>size | Median<br>Value | Range |  | Environment | Reference |
| --- | --- | --- | --- | --- | --- | --- |
|  |  |  | Lower<br>value | Larger<br>value |  |  |
| Clade I | 3 | 12,822,270 | 10,110,043 | 16,231,765 | Sediment | This work |
| Clade II |  |  |  |  |  |  |
| Sub-clade A | 4 | 55,938 | 34,079 | 392,104 | Sediment | This work |
| Sub-clade D | 6 | 20,849 | 2,795 | 218,603 | Water-Sediment | This work |
| Sub-clade G | 4 | 29,791 | 6,174 | 226,658 | Water-Sediment | This work |
| Clade III | 4 | 15,018,880 | 8,970,283 | 22,432,331 | Water-Sediment | This work |
| Clade IV | 4 | 383,067 | 345,564 | 427,557 | Sediment | This work |
| Clade V | 4 | 9,594,874 | 5,894,074 | 12,914,770 | Sediment | This work |
| Clade VI | 3 | 4,141,870 | 2,582,483 | 10,645,019 | Sediment | This work |
| *H. pylori* |  | 39,665,437 | - | - | - | [108] |
| *S. enterica* |  | 348,991,354 | - | - | - | [108] |
| *E. coli* |  | 179,600,000 | - | - | - | [108] |
| *H. sapiens* |  | 20,348 | - | - | - | [108] |
| *A. thaliana* |  | 266,769 | - | - | - | [59] |
| *C. elegans* |  | 3,998,701 | - | - | - | [109] |
| *T. brucei* |  | 5,332,244 | - | - | - | [108] |

Summary table of effective population sizes of CCB Clades and prokaryotic and eukaryotic references. First column shows the names of the CCB Clades and reference strains used for the calculus, second column represents the number of strains within each group, followed by the median Ne value estimated and the range. Last two columns display the isolation environment and the reference.

**Table 5.** GO terms enriched estimated with TopGO [45], regarding the gene families with signals of positive selection.

| GO.ID | Term | Annotated | Significant | Expected | Fisher test with Bonferroni |
| --- | --- | --- | --- | --- | --- |
| GO:0000902 | cell morphogenesis | 398 | 67 | 26.93 | 0.00020748 |
| GO:0009234 | menaquinone biosynthetic process | 240 | 38 | 16.24 | 0.00150024 |
| GO:0009245 | lipid A biosynthetic process | 240 | 38 | 16.24 | 0.00150024 |
| GO:0008360 | regulation of cell shape | 244 | 37 | 16.51 | 0.0059052 |
| GO:0007156 | homophilic cell adhesion via plasma membrane adhesion molecules | 13 | 7 | 0.88 | 0.0122892 |
| GO:0006304 | DNA modification | 295 | 41 | 19.96 | 0.01596 |
| GO:0009058 | biosynthetic process | 26775 | 1675 | 1811.62 | 0.017556 |

First two columns show the enriched GO IDs and its name, third column the number of annotated genes, fourth and fifth column the number of significant genes and the expected, last column shows the significance corrected with Bonferroni.

### Pan-genome analyses of CCB Vibrionaceae and lineages description

The pan-genome analysis of 39 CCB *Vibrio* spp., 3 CCB *Photobacterium* spp., and 5 *Vibrio* spp. references strains involved a total of 20,121 orthologous gene families. The genes that were present in at least 95% of the genomes conformed the core genome, including reference genomes, composed by 1,254 gene families. The accessory genome is far more substantial, consisting of a total of 14,072 genes families that were found in at least two of the obtained genomes. The rest 4,795 genes families were strain-specific.

Using the core genes families, we reconstructed a lineage phylogeny (Figure 2). In the core phylogeny we found seven lineages, of which six of them were previously identified in the 16s rRNA gene tree, and one was represented by a unique strain of marine *V. furnissii* sp. Nov. 4 stran (NCTC 11218) [39]. Reference strain *V. anguillarum* 775, isolated from a Coho salmon [40] clusters within the large generalist Clade II, while reference strain *V. metschnikovii* CP 69-14, which was isolated in marine systems, is basal to Clade III. Basal to Clade VI are reference *V. parahaemoliticus* BB22OP, a pre-pandemic strain [41] associated with seafood-borne gastroenteritis in humans and *V. alginolyticus* NBRC 15630 = ATCC 17749, an aquatic organism that can cause bacteremia. Clades IV and V are likely to be exclusive to CCB, given that there is no closely related strain sequenced on databases. Finally, Clade I is related to *Photobacterium* spp. (Figure 2).

From the six clades identified, only Clade II presented an open pan-genome as suggested by the Heaps law analysis [42] (alpha= 0.7913). The rest of the clades displayed closed pan-genome patterns (i.e., alpha values >1.0; Table 1). We performed random sub-samplings of genomes per clade to verify the effect of sample size and we re-calculate alpha values of each clade with the minimum sample size, and in all cases, we found the same results of closed pan-genomes for the specialist clades and an open pan-genome for generalist clade (Additional file 1: Figure 3).

**Figure 3.**
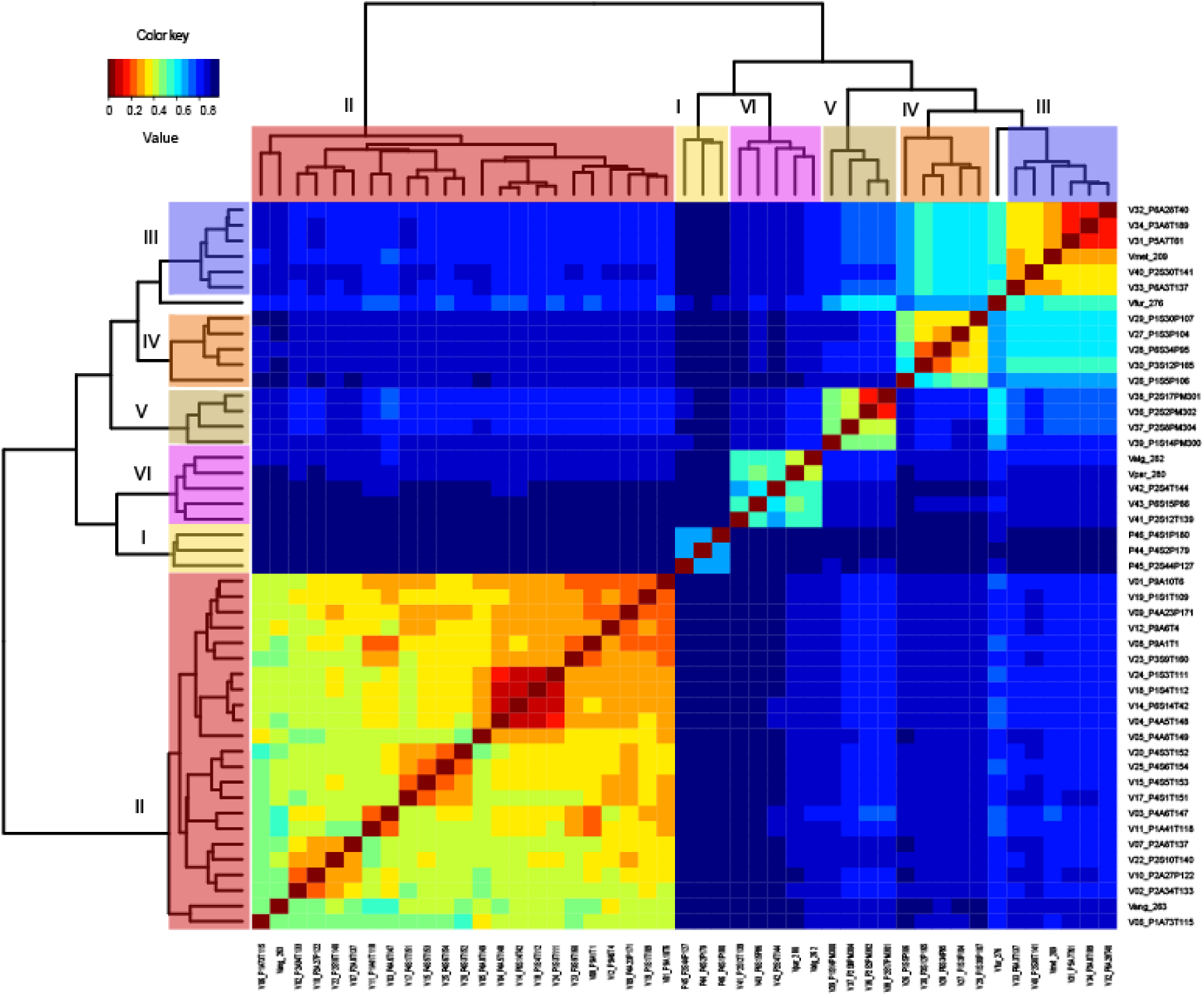
Patterns of recombination events among isolated strains. Heatmap of the frequency of recombination events among different strains; red colors indicates more recombination events within strains while blue events indicate few recombination events. Distances were estimated with the Jaccard dissimilarity index.

### Genetic diversity and recombination estimates

General estimators of genetic diversity were obtained for each clade and Sub-clade (Table 2). We found that nucleotide diversity values for Clades III, IV, and V were the lowest within sample, ranging from 2.86E-05 to 0.0051, while Clades I, II, and VI had higher levels of genetic variation, in the range of 0.011 to 0.046. When estimating the nucleotide diversity for Sub-clades belonging to Clade II (described below, see Additional file 1: Figure 4), we found lower values in the range of 1.61E-06 to 5.47E-06. This same pattern was observed for the θw values (Table 2). Due to the number of individuals we could not obtain Tajima’s *D* estimator for Clades I and VI. For the rest of the clades Tajima’s *D* values were negative, except for Clade II that had positive values.

**Figure 4.**
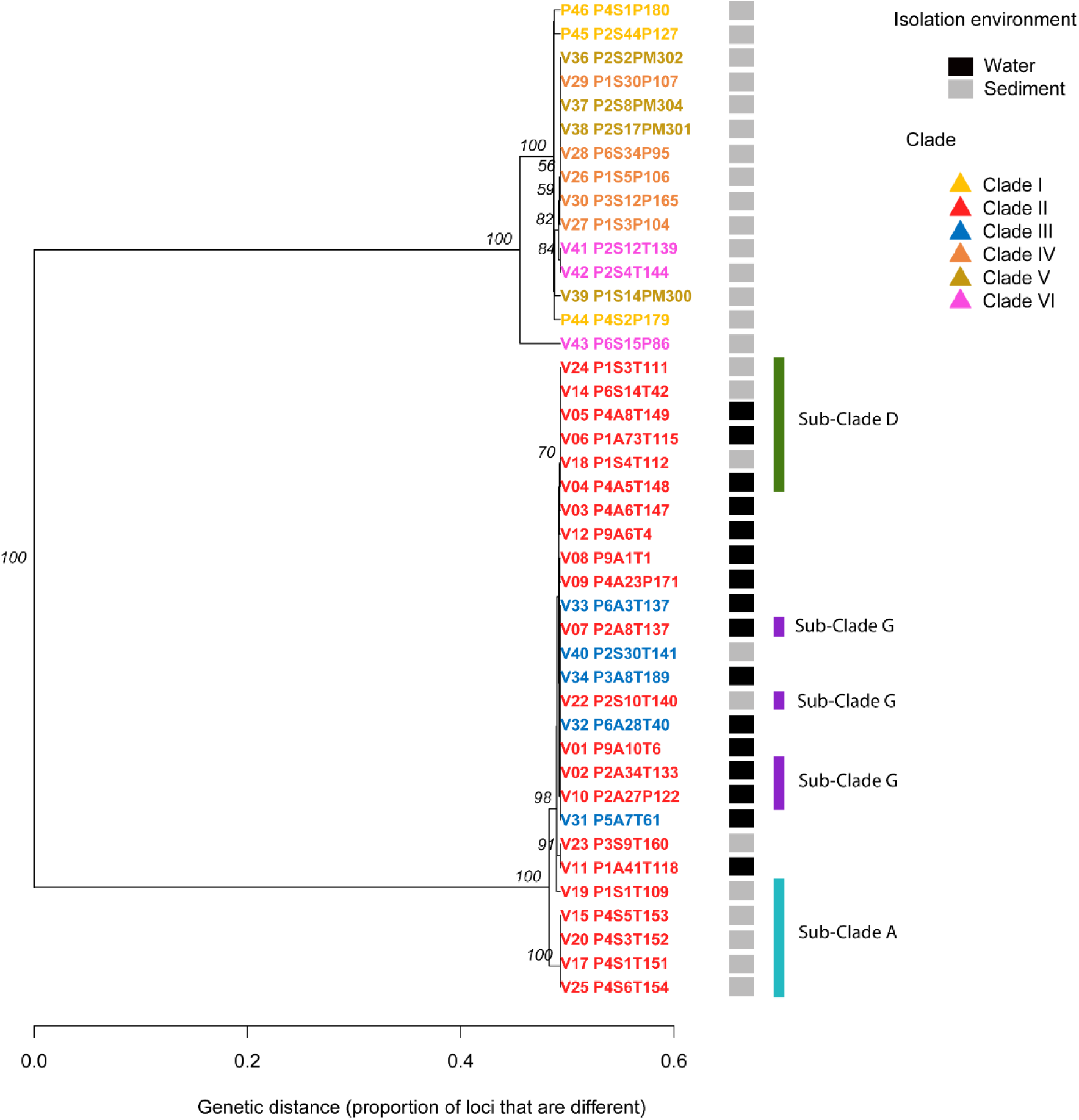
UPGMA of the 598 SNPs associated with the isolation environment. Tip colors represent clade membership, for Clade II, Sub-clades are also indicated. Squares represent the isolation environment. Distances were calculated with the bitwise distance function of poppr v2.8.1.

Since most lineages present a closed pan-genome, we tracked the footprints of recombination by using two different approaches. The first approach consisted of assessing the recombination in each ortholog group. The second involved the identification of recombination signals based on a whole genome alignment. With the first approach, we found that from the 15,380 ortholog clusters analyzed, only the 11% (1,759) showed significant signal of recombination (Additional file 2: Table 6). These recombination events occurred more frequently among isolates of the same environment and pond, suggesting reproductive isolation associated to an environmental variable (Figure 3). However and despite we considered in our calculations the pan-genome size, number of strains per clade and branch length, it is also true that most clades are conformed by only isolates of water or sediment. Therefore, we propose that the frequency of recombination events is mostly restricted to occur within clades (Figure 3; Additional file 1: Figure 5).

**Table 6.**
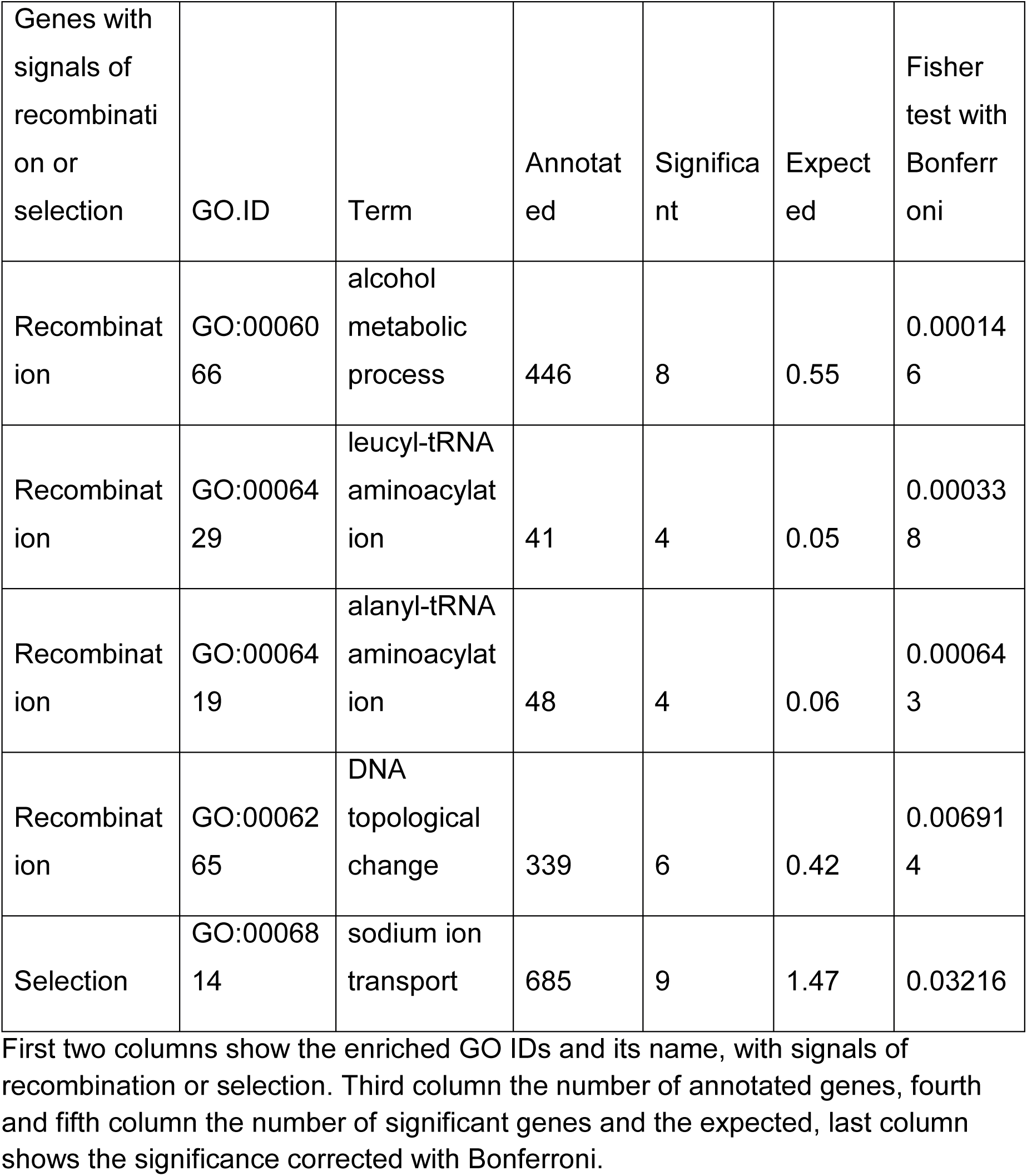
GO terms enriched in the genes found to have an association with the isolation environment (water or sediment).

In the case of the generalist Clade II, we found sub-structure. Using Nei’s genetic distances, we identified ten genetic groups (that we will call Sub-clades therefore) with distances greater than 0.001. The discriminant function shows the same structure as the Nei distances, reflecting a broader relationship between Sub-clades A, D, F and G and B with C and E. Meanwhile, H, I and J Sub-clades had dissimilar sub-structures (Additional file 1: Figure 4). Since only three of the Sub-clades contained more than two isolates, further analyses were just performed with the larger Sub-clades (A, D and G).

Following the second approach, we evaluated the impact of homologous recombination and mutation within lineages estimating *r/m* using the clonal frame software [43]. This measure reflects the ratio of probabilities that a given polymorphism is explained by either recombination (*r*) or by mutation (*m*). Clade VI displayed the lowest *r/m* values=0.0052, while Clade I (i.e., *Photobacterium* spp.) had the highest value in our dataset, *r/m* = 2.72 (Table 3). We also performed the same analysis on *V. parahaemolyticus, V. ordalii, V. anguillarum*, and *P. leiognathi* reference genomes, all isolated from marine environments. For the marine samples, *r/m* estimates were within the range of CCB strains, except *V. anguillarum*, which had the highest values (Table 3). This analysis also shows that some recombination events are shared with *Vibrio* spp. references strains (Additional file 1: Figure 6) supporting the hypothesis of ancient origin of these recombination events even though more recent recombination events were detected only among CCB strains. This indicates that homologous recombination is a constant source (albeit relatively infrequent) of polymorphism in the analyzed strains.

### Estimates of effective population sizes

Using a simulation approach with the Fastsimcoal2 program [44] we estimated the posterior distribution of the effective population size (*N_e_*) of each of the six clades. We found large population sizes (Table 4) ranging from millions in the specialist Clades I (*N_e_* = 12,822,270), III (*N_e_* = 15,018,880), V (*N_e_* = 9,594,874) to intermediate in the range of thousands in the Clades IV (*N_e_* = 383,067) and VI *(N_e_* =141,870), and to far smaller in the Sub-clades of the locality common generalist Clade II (i.e., Sub-clade A *N_e_* = 55,938; Sub-clade D *N_e_* = 20,849; Sub-clade G *N_e_* = 29,791) reinforcing the idea of recent diversification in these Sub-clades.

### Selection analyses

FUBAR uses a codon-based model of evolution that allows the identification of evolving sites under positive or purifying selection in protein-coding genes through a Markov chain Monte Carlo (MCMC) routine. From a total of 15,380 ortholog clusters analyzed, only 367 (2.3 %) had a significant signal of positive selection according to FUBAR results. Of these ortholog gene families, 297 belonged to the flexible genome, while 70 are part of the core genome. However, when we considered the universe of ortholog genes that conform the flexible genome (14,072), only 2.1% of the flexible genome had signals of positive selection, while in the core genome, composed by 1,254 genes, 5.6% of the genes are positive selected (Additional file 2: Table 7). A GO enriched analysis was performed in order to identify those biological functions overrepresented given those ortholog clusters with positive selection. Seven Gene Ontology (GO) terms were enriched within these families. (Table 5). Moreover, based on a whole genome alignment, we obtained 38,533 SNPs variants, from which 26,663 were bi-allelic characters that were used in an UPGMA analysis of genetic distances. This analysis produced the same clustering as the core genome phylogeny (Figure 2). As well, with this SNPs we performed a membership probability test, which show that all the isolated had the same probability of been isolated from any pond and environment (Additional file 1: Figure 7).

We found on average 2,473 private (unique) SNPs for each one of the nine ponds, 33,655 private SNPs for water or sediment environments, and 29,141, private SNPs for each of the six clades. This abundance of private SNPs suggests an effect of the environment, either by local adaptation (selection) or by genetic drift (low effective sizes or little or no gene flow).

We removed the SNPs with a minor allele frequency < 0.05 (771 SNPs removed) and we kept the alleles that were found in at least three individuals, for a total of 25,892 SNPs. Within those SNPs we detected a total of 598 SNPs with an association to the sediment environment. A UPGMA analysis of these 598 SNPs was performed in order to infer the similarity between samples (Figure 4) finding most of the clusters previously observed with the core genome phylogeny (Figure 2), except Clade III, which appears inside Clade II. Moreover, the mixed isolates of Clade III fall among the Sub-clade G of Clade II, most of them were isolated from water environment, as well as members of Clade III (Figure 4), suggesting a preference for diluted, unstructured environments.

To analyze the distribution of the SNPs, we mapped the above detected 598 SNPs to their positions in the genome alignment from where they were obtained, moving in 1 Kb windows. A total of 144 genomic regions containing SNPs were inspected, and we found 237 ortholog gene families in these regions. From these ortholog gene families, only 24 showed recombination signals, while 18 had selection signals (Additional file 2: Table 8). Within those SNPs we performed a test for GO-term enrichment with TopGO [45]. From the 24 ortholog genes families with recombination signals, we detected four enriched GO, while we found only one enriched GO-term in the 18 ortholog gene families with selection signals (Table 6).

Besides those analyses, based on pan-genome information, we looked for specific coding sequences that could be private (unique) to a specific pond, environment, or clade. There were no specific genes associated with a particular environment or pond, but we did identify ortholog gene clusters exclusive per clade. From Clades I to VI, we observed 1280, 10, 72, 23, 72, and no exclusive ortholog gene families, respectively. For each clade with exclusive ortholog gene families, we looked for enriched GO terms. On Clade I the term related with bacteriocin immunity was enriched; Clade II were enriched with terms associated to siderophore transport; in Clade III the category related to the biosynthesis of lipopolysaccharides was enriched; and on Clades IV and V there were enriched terms related to tRNA biosynthesis (Additional file 1: Table 9).

## Discussion

In this study we performed comparative genomic analyses to understand how evolutionary forces shaped the pan-genome of 42 Vibrionaceae strains isolated from CCB, where environmental filtering is believed to increase local adaptation due to extreme stoichiometric bias [27]. In our study we described how a natural perturbation lead to a temporal balanced stoichiometry, allowing six lineages of Vibrionaceae to prosper under a “feast-famine” cycle. Most of these lineages present large population sizes as well as recombination rates comparable to their oceanic counterparts. However, their pan-genomes remained closed probably due to selection purging HGT events external to each clade where genetic isolation has maintained clade specific selective events. Clade II is the exception, this large clade shows an open pan-genome with evidence of substructure with small effective sizes suggesting early stages of diversification.

### Ecology and microbial diversity in CCB

During the past 20 years, one of the main questions surrounding CCB bacterial hyper-diversity has been related to the roles of ecology and evolution promoting and maintaining its remarkable microbial diversity [27, 46]. According to Souza *et al*. “lost world” hypothesis, the extreme unbalanced stoichiometry (i.e., very low P availability) of CCB not only keeps the “ancestral niche” of many bacterial lineages, but also works as a semipermeable barrier to migration, restricting migration and keeping these ancient bacterial lineages alive and thriving in CCB [27]. As a result of these ecological and evolutionary conditions, CCB lineages are generally clonal [28–30] displaying an ancient marine ancestry [27, 32, 47]. Paradoxically, this extremely unbalanced stoichiometry seems to be in part the reason behind CCB high microbial endemicity and local differentiation: “No food, no sex, no travel” [27, 31, 32], allowing for local adaptation and broad differentiation between sites.

In this study we explored the evolutionary dynamics after a natural perturbation (in this case a flood) changed the ecological condition in CCB in a particular site (Pozas Rojas), generating a temporarily more “balanced” stoichiometric proportions (i.e., N:P 20:1). We know by meteorological data that similar floods occur at CCB sporadically, due to the low incidence of intense storms (i.e., three since 1940 [48]). The flood moved to this low land a large amount of debris that with time, generated an increase in nutrients, in particular phosphorus that opened opportunities for the “rare biosphere”, represented by standing bacterial lineages usually found at very low proportions, like the rare members of Vibrionaceae that normally are not common at standard low nutrient conditions [49–51]. Given this change in resources, we proposed two hypotheses when we started this study: Vibrionaceae from CCB would show as their ocean counterparts, an open pan-genome, showing high levels of recombination and genetic variation, as well as a high N_e_. Alternatively, due to local adaptation in each lineage of CCB, Vibrionaceae would display closed pan-genomes, and a strong genetic structure, generated by high clonality and low genetic variation probably related to periodic selection and small effective population sizes among lower levels of genetic variation.

### Vibrionaceae in CCB

In a previous study at Pozas Rojas using both cultivated strains and metagenomic data, Bonilla-Rosso *et al*. found that *Vibrio* spp. was either very rare or absent [49]. In their study, the authors found mostly Pseudomonads among the cultivated strains [49]. This result was confirmed with metagenomics, where Pseudomonadales, Burkholderiales, and Bacillales represented 50% of the metagenome reads. As a result of this previous knowledge, in the 2013 sampling, we first used PIA media to analyze the effect of the 2010 flood in the previously abundant genera, however, we found that this lineage was replaced in the cultures by *Vibrio* spp. In other words, the increased levels of nutrients and the perturbation reduced the abundances of *Pseudomonas* and related genera in CCB. This effect was corroborated later in another system in CCB (Churince) with a nutrient enrichment experiment [50, 52]. Among the analyzed genomes, we found two clades of Vibrionaceae, Clades III and IV, that had not been isolated previously and could be endemic to the basin.

### Recombination, pan-genomes, and selection in Vibrionaceae

Diversity measures, π and θw showed lower diversity than cosmopolitan *E. coli* [53], nevertheless, for Clades I, II and VI, those values are comparable to the ones observed in pathogenic *Vibrio* spp. [54, 55] suggesting similar demographic dynamics. Tajima’s D was in most cases negative, except for Clade II, but none of the values were statistically significant. This could suggests bottlenecks in the process of diversification explaining the extremely low effective population size and diversity in those Sub-clades. Negative values of Tajima’s D suggest high content of rare alleles, which is in agreement with the private allele test we performed [56]. In the same way, it could be the result of selective sweeps or recent demographic expansion as a result of the new nutrient conditions (feast).

This study corroborates the importance of recombination in Vibrionaceae, supporting the recombinant nature of the genomes in the family [33, 34]. Elevated recombination rates are maintained in all the lineages from Pozas Rojas, supporting our first hypothesis that *Vibrio* spp. from CCB would have similar evolutionary process and genetic structure than marine lineages. Further scrutiny revealed an unexpected result: even if recombination rate is similar to their oceanic counterparts, homologous recombination and selection apparently maintain the adaptation to the local environment. Even in Clade II, recombination is more abundant among related strains, suggesting that this clade is in an actively diversifying process, allowing their different Sub-clades to adapt to different environments within CCB, as it is the case of aquatic Sub-clade G that shares similar SNPs under selection than aquatic Clade III.

We believe that the natural disturbance at Pozas Rojas generated by an increase in nutrient availability relaxed selection against HGT. Nevertheless recombination is kept within close lineages resulting in large effective population sizes and a closed pan-genome in most of the lineages, allowing selection to act in response to environmental pressures [57–59]. The closed pan-genome of these lineages contrast to what has been reported in oceanic *Vibrio* spp. where populations sizes are large and pan-genomes are kept open due to HGT [60]. Even though Clade II is the only one with an open pan-genome, its internal substructure suggest a recent process of diversification where each of its Sub-clades show again a closed pan-genome, with smaller *N_e_* and low genetic diversity.

### Selection and adaptation in Pozas Rojas

We found 367 gene families that have signals of positive selection, most of them regarding the whole group of ortholog genes found in the core genome (2.05% of the flexible genome and 5% of the core genome; Additional file 2: Table 9). This result suggests that selection purges the genes that are in the flexible genome, closing the pan-genomes. Among the detected genes with selection signals, seven functional GO terms were enriched, one of them was the term GO:0007156, which is associated with cell-cell adhesion; within this category, most of the genes annotated were related to cadherin domains that have been associated to biofilm formation [61]. In natural environments, biofilm formation allows bacteria to cope with environmental changes, protects the cell, provides mechanical stability, and provides cellular adhesion with other cells or with surfaces. It has been observed that biofilm formation is a persistent characteristic among bacteria from CCB in both water and sediment, and also under different nutrient conditions [52].

When we performed a genome-wide association study (GWAS) test to analyze the association of the SNPs to either water or sediment environment, we identified 598 SNPs related to sediment. The UPGMA analysis showed a similar clustering pattern as the core genome (Figure 4), suggesting a clade effect. However, Cluster III grouped among the Sub-clade G of Clade II, and most of the isolates of this Sub-clade as well as Clade III were isolated from the water environment. One possibility is that these SNPs are important to the adaptation to non-structured environments such as water. Some of the genes associated to these SNPs presented signals of recombination and selection. One of the functional enrichment GO terms within these genes was the GO:0006814, which is involved in sodium transport; some of the genes annotated within this category were the bacterial Na+/H+ antiporter B (NhaB) that has been suggested to play a role in the adaptation of halophilic and haloalkaliphilic proteobacteria to marine habitats [62]. This gene has also been found to play a role in homeostasis in *Vibrio* spp. [63]. Our data suggest that there is a selective pressure over some clades regarding the water environment.

When we analyzed unique genes for each clade disregarding the isolation environment, in the case of Clade I we found the term GO:0030153 enriched, which is related to bacteriocin immunity. However, antibiotic resistance associated genes did not show particular signals of selection, suggesting that overall there is no ongoing selective pressure for defense. In the large generalist Clade II, we found three GO terms enriched, two of them related to cell wall structure while the third is related to siderophore transport, a group of genes that were rare in the previous metagenomic analysis of the same site [35]. In the case of Clade III, the enriched GO term is related to lipopolysaccharide biosynthesis. Meanwhile, in Clade IV, we identified six enriched GO terms, where most of them were related to transport and signal transduction. Finally, for Clade V, we identified four terms enriched mostly related to transport. These results suggest that distinct clades are indeed responding to their environment in different ways reinforcing the idea of genetic isolation as a way to preserve local adaptation (Additional file 2: Table 8).

### Perspectives and conclusions

At CCB, most of the environments present an extremely low phosphorus concentration, a factor that acts as an effective migration barrier maintaining conditions of the ancient sea as well as ancestral microbial diversity [27]. However, due to natural perturbation, we had the opportunity to observe in Pozas Rojas what happens when that nutrimental barrier is lifted temporarily. Apparently, rare biosphere strains that normally had a hard time surviving low P conditions can follow a feast-famine cycles and have population expansion when the P availability is less limiting.

In order to understand the other dimensions of local adaptation, further sampling of *Vibrio* spp. in CCB is needed. Unfortunately, this extraordinary oasis is disappearing, given the loss of more than 95% of CCB wetlands due to groundwater overexploitation by agriculture [27, 47,51, 64].

## Methods

### Site description

We analyzed bacterial isolates from sediment and water of nine ponds in the Pozas Rojas area of CCB (Figure 1). This site is composed of several small ponds (locally called *pozas*) that surround a larger pond in the system of Los Hundidos [30, 35]. These small ponds become hypersaline in summer [30], and used to have the highest stoichiometric unbalance (i.e., lowest P concentration) reported in CCB (C:N:P 15820:157:1)[35]. The ponds have seasonal high fluctuations in temperature (around 1 °C in winter to up to 60 °C in some summer moments in some cases)[35] and are small but permanent, separated from each other by ca. 9 meters or more, along an arch around the larger pond. However, the Pozas Rojas were flooded by hurricane Alex during summer 2010, merging most of the small ponds into a single large pond, until autumn 2011, when the water receded, leaving the moon shaped array of small red ponds at the same place (Figure 1).

### Sample collection and strains isolation

We collected water and sediment samples in duplicate from nine ponds located in Pozas Rojas, Los Hundidos, CCB, during March 2013 and stored them at 4 °C until processing. Sediment was collected for nutrient analysis in 50 ml Falcon tubes and covered with aluminum foil before storage. Water was collected for nutrient quantification in 1 liter volumes and stored in the dark at 4 °C. Chemical analyses were performed at the Instituto de Investigaciones en Ecosistemas y Sustentabilidad, UNAM, in Morelia, Mexico. Cultivable strains from both sediment and water were isolated in PIA (*Pseudomonas* isolation agar) and TCBS (Thiosulfate Citrate Bile Sucrose Agar) as previously described [52, 65], obtaining a total of 174 isolates, being 88 isolates from sediment and 86 from water.

### Environmental variables measurement

For nutrient quantification, sediment samples were dried, and water samples were filtered through a Millipore 0.42 µm filter. Total carbon (TC) and inorganic carbon (IC) were determined by combustion and colorimetric detection [66] using a total carbon analyzer (UIC model CM5012, Chicago, USA). Total organic carbon (TOC) was calculated as the difference between TC and IC. For total N (TN) and total P (TP) determination, samples were acid digested with H_2_SO_4_, H_2_O_2_, K_2_SO_4_ and CuSO_4_ at 360°C. Soil N was determined by the macro-Kjeldahl method [67], while P was determined by the molybdate colorimetric method following ascorbic acid reduction [68]. The N and P forms analyzed were determined colorimetrically in a Bran-Luebbe Auto analyzer 3 (Norderstedt, Germany).

### DNA Extraction and PCR Amplification of 16S rRNA

For the 174 isolates obtained, DNA extraction was performed as described by Aljanabi and Martinez (1997) [69]. 16S rRNA genes were amplified using universal primers 27F (5′-AGA GTT TGA TCC TGG CTC AG-3′) and 1492R (5′-GGT TAC CTT GTT ACG ACT T-3′) [70]. All reactions were carried out in an Applied Biosystems Veriti 96 Well Thermal cycler (California, USA) using an Amplificasa DNA polymerase (BioTecMol, Mexico) with the following program: 94°C for 5 min, followed by 30 cycles consisting of 94°C for 1 min, 50°C for 30 s, 72°C for 1 min and 72°C for 5 min. Polymerase chain reaction (PCR) amplification products were electrophoresed on 1% agarose gels. Sanger sequencing was performed at the University of Washington High-Throughput Genomics Center.

### Phylogenetic analysis of 16S rRNA sequences

The first 700 bps of the 16S rRNA gene, were aligned with Clustalw [71] and quality control was performed with Mothur [72]. Genera level identification was made using the classifier tool [73] from the Ribosomal Database Project (RDP) Release 11.4 [74] (Additional file 1: Table 3). Blastn searches were performed against Refseq database from NCBI to select reference sequences. A total of 101 sequences were identified as members of the Vibrionaceae family, 41 were isolates from water and 60 from sediment. These isolates were used in subsequent analyses. A maximum likelihood phylogenetic reconstruction was obtained with PhyML version 3.0 [75], using the HKY+I+G substitution model estimated with jModelTest 2 [76]. The degree of support for the branches was determined with 1,000 bootstrap iterations.

### Environmental association of phylogroups

To test whether the community of cultivable strains was structured based on its isolation environment (i.e., water or sediment), we performed an AdaptML analysis [37], including our 101 isolates belonging to Vibrionaceae and an *Halomonas* spp. strain as an out-group. Three categorical environmental variables were tested, including pond of isolation, high and low nutrient concentrations, and the two sampled environments (water or sediment).

### Genome sequencing, assembly, and annotation

For whole-genome sequencing, we selected from the AdaptML analysis 39 *Vibrio* spp. isolates, 23 isolated from sediment and 16 from water, plus 3 isolates of *Photobacterium* spp. (a lineage closely related to the *Vibrio* spp. genus) isolated from sediment. DNA extractions were performed with the DNeasy Blood and Tissue kit (Qiagen).

Sequencing was performed with Illumina MiSeq 2×250 technology, with insert libraries of 650 bps and an expected coverage of ca.10x per genome. At first, we planned an assembly strategy using a genome reference; for this reason, the strain V15_P4S5T153 had a second library that was designed using the Jr 454 Roche technology, in order to reduce sequencing bias and get higher coverage. However, due to divergence among genomes, we performed *de novo* assemblies for all genomes. All sequencing was performed at the Laboratorio Nacional de Genómica para la Biodiversidad (LANGEBIO), México.

The quality of raw reads was analyzed using FASTQC software (http://www.bioinformatics.babraham.ac.uk/projects/fastqc/). A minimum quality value of 25 was set, and low-quality sequences were removed with fastq_quality_filter from the FASTX-Toolkit (http://hannonlab.cshl.edu/fastx_toolkit/index.html). Adapter sequences were identified, removed and paired-end reads were merged using SeqPrep (https://github.com/jstjohn/SeqPrep). *De novo* assemblies were performed with Newbler (Roche/ 454 Life Sciences) using both single-end and merged reads.

For scaffolding process, we used SSPACE [77], gaps were closed using GapFiller [78] and final error correction was performed with iCORN [79] (Additional file 2: Table 10). Coding sequences were inferred with Prodigal 2.0 [80] implemented in PROKKA software [81]. InterProScan 5 allowed annotation [82] with the databases enabled by default. Genome completeness was assessed with BUSCO using the Gamma-proteobacteria database [38].

### Pan-genome analyses

The 42 genomes from CCB where compared with genomes of 5 reference *Vibrio* spp. strains: *Vibrio alginolyticus NBRC 15630 = ATCC 17749*, *V. anguillarum 775*, *V. furnissii NCTC 11218*, *V. parahaemolyticus BB22OP* and *V. metschnikovii CIP 69 14* (Additional file 1: Tables 4, 5; Additional file 2: Tables 10). Ortholog gene families were predicted from all 47 genomes using the DeNoGAP comparative genomics pipeline [83]. To minimize false positive prediction of orthologs, we assigned *Photobacterium* spp. genomes as outgroup. The completely sequenced genome of *V. anguillarum* strain 775 was used as seed reference.

We estimated the core genome based on presence and absence of gene families across the genomes. If the genes were present in all strains, the orthologs were classified as *core*, while genes were classified as *accessory* when present in more than one strain but not in all of them, and *unique* genes when it was present only in a single strain. Since most of the genomes in our dataset are not completely sequenced, we designated core ortholog families as those present in at least 95% of the genomes, to avoid the impact of missing genes due to sequencing or assembly artifacts.

The package Micropan [84] within R v.3.4 (R Core Team) [85] was used to infer the open or closed nature of each pan-genome dataset, following the heaps law proposed by Tettelin *et al*. [42]. The Heaps law model is fitted to the number of new gene clusters observed when genomes are ordered randomly. The model has two parameters: an intercept, and a decay parameter called alpha. If alpha is higher than 1.0 the pan-genome is considered closed, if alpha is lower than 1.0 it is considered open. Additionally, a random sub-sampling for each clade was made, taking three genomes and calculating the alpha value for each group of three genomes. A total of 1,000 independent sub-sampling events were made for each clade.

Core proteins were aligned using Kalign [86] to infer the phylogenetic relationship between the samples. The resulting alignments of individual ortholog families were concatenated using a custom Perl script. With these concatenated core genes, a maximum likelihood phylogenetic tree was constructed using the FastTree program [87].

### Recombination analyses

Of the total ortholog families in the *Vibrio* spp. pan-genome, we only used the ortholog families found in at least three genomes for the recombination analyses. Genetic recombination was examined on each CDS alignment by using inference of pairwise recombination sites, obtained with GENECONV [88] and by the identification of putative recombinant sequences through breakpoints using GARD [89].

Based on the number of recombination events, we estimated the events shared among isolates of the same pond and environment, among isolates of the different pond and environment, among isolates of the same pond and different environment and among isolates of different pond and environment. For this, we normalized the data by pan-genome size, number of strains and branch length. Given that the large generalist Clade II presented a clear sub-structure, we did a separated analysis for the shorter branches within Clade II (Additional file 1: Figure 4).

To assess the impact of homologous recombination, we analyzed the substitution pattern using two different algorithms, Gubbins [90] and ClonalFrameML [43]. A whole-genome alignment for the 47 analyzed genomes was performed with MAUVE [91]. The resulting alignment was used as input for Gubbins [90] using RAxML [92] and default parameters. Additionally, whole genome alignments were performed for each clade, excluding references, with the progressive MAUVE algorithm [91]. We calculated the R/theta ratio, nu and delta [43] for each sample and for 100 bootstrapped replicates.

### Genetic structure of Clade II

Recombination analyses showed that in Clade II there are internal groups with higher internal recombination, so we decided to further investigate the structure within Clade II. For clustering analyses, we used Nei’s genetic distance [93] and neighbor joining. Genomes with distance less than 0.001 were grouped and tested with a discriminant analysis of principal components of the genetic variation, using the adegenet library in R [94]. For this study, we used 20 principal components and 3 discriminant functions.

### Selection analyses

We used FUBAR [95] to identify signatures of positive selection among ortholog gene families found in at least three genomes. We accounted for recombination breakpoints in the ortholog families, while calculating positively selected sites based on GARD results [89]. We considered any site to be positively selected if it showed P-value ≤ 0.05. We also conducted a Gene Ontology (GO) enrichment analysis using topGO [45] to find overrepresented biological functions in this set of genes.

### Effective population size estimation

We followed a simulation approach to estimate the posterior distribution of the effective population size (*N_e_*) of each of the six clades. According to the previous clustering and recombination analysis, for Clades I, III, IV, V and VI we simulated a single population, while for Clade II we simulated three sub-populations that diverged from an ancestral population.

Simulations were performed using Fastsimcoal2 [44, 96]. For each clade, we simulated DNA sequences having a similar length equal to the number of nucleotides in the given clade, as well as a sample size equal to the number of sequences sampled for each clade. We assumed no recombination within the genome, and used the *Escherichia coli* mutation rate of 2.2×10^−10^ mutations per nucleotide per generation [97]. We ran between two and four simulations for each clade. For the initial runs, we generated 100,000 replicates extracting *N_e_* values from a prior log-uniform distribution that ranged from 100,000 to 20,000,000 individuals. For Clade II, we also estimated the age of divergence of each Sub-clade, by setting the prior distribution of time ranging from 1,000-4,000,000 generations. After a first run, we narrowed the prior ranges based on those simulations that had similar summary statistics compared to the observed data and performed another 100,000 simulations using the narrowed priors.

To compare the previously simulated and observed data based on summary statistics, we used the ape [98] and pegas [99] libraries in R to estimate the number of polymorphic sites and the Tajima’s *D* based on the entire genomes. Tajima’s *D* is commonly used to estimate demographic changes in populations [100, 101]. Also, we obtained 1,000 sliding windows frames to estimate the Tajima’s *D* along the genomes, as well as the mean and standard deviation of Tajima’s *D*. Tajima’s *D*, π, and Watterson’s theta (θw) were estimated for each clade as well as for Sub-clades A, B and G. Since clades I and VI had three sequences and it was not possible to obtain Tajima’s *D*, we did 1,000 replicates in which we subsampled with replacement 10 sequences. For each replicate, we calculated Tajima’s *D* and we obtained as the proximate value the median estimated across the 1,000 replicates.

Based on the summary statistics, we used the abc function in the ABC package [102] in R to calculate the distribution of the *N_e_* parameter based on a 0.05 % threshold distance between the simulated and observed data. For each clade, we report the median and the 95% interval confidence of *N_e_*. For Clade II, we further reported the average and 95% interval confidence of the number of generations since each Sub-clade diverged from an ancestral clade.

### Association between genotypes and environmental variables

We evaluated whether the genetic variation within the Vibrionaceae genomes could be explained by particular adaptations to the environment (water or sediment). We used progressiveMauve [91] to perform a global multiple alignment between the assembled genomes. We extracted the variant sites within the alignment and exported them as SNPs using snp-sites [103].

We obtained 38,533 SNPs, which we used to search for private alleles using Poppr [104]. Afterwards, we obtained a subset of 25,892 SNPs by filtering biallelic sites with minor allele frequencies > 0.05. We used PLINK [105] to perform a GWAS to detect possible associations between our SNP set and either the water or sediment environments. We conducted Fisher exact tests and regarded as significant all SNPs whose associations had *p*-values < 0.01 after Bonferroni corrections. These analyses may be informative even considering these sampling differences [106, 107].

To test whether these associations could be explained by convergent evolution rather than by common ancestry, we compared an UPGMA tree reconstructed from the total set of SNPs from an UPGMA tree using only the SNPs that were significantly associated to the environment. We analyzed the distribution of the SNPs within the genomes to find the genes associated to those SNPs.

We mapped the SNPs positions in the genome alignment moving by 1 Kb windows; this window size was selected considering the average bacterial gene size and retrieved all the associated genes. We conducted a Gene Ontology (GO) enrichment analysis using topGO [45] to find overrepresented biological functions in this set of genes.

## Supporting information

AdditionalFile1

AdditionalFile2

## Availability of data and materials

The datasets generated and analysed during the current study are available in the genome assembly project BioProject: PRJNA361510; PRJNA361511. The resulting InterProScan annotation files, CDS fasta files and the predicted protein fasta files for all taxa are available at Dryad. As by the politics of Dryad, the data will be available once the manuscript is accepted.

## Author Contributions

MV-R-L design the sampling, obtained the biological material, analyzed the data, prepared figures and tables, and wrote the paper. GYP-S analyzed the data and participated in all stages of writing. JA-L, ST, ES, and JB-R analyzed the data. EI-L analyzed the data and provided computing facilities. DS-G provided computing facilities and contributed substantially to the analysis and discussion of the data. LEE made contributions for the design, analysis, discussion of the data and writing. V-S conceived, designed the study and the analyses, managed the obtaining financial resources and participated in all stages of writing.

## Acknowledgments

We thank Felipe García-Oliva and Rodrigo Velázquez-Durán at the Instituto de Investigaciones en Ecosistemas y Sustentabilidad, UNAM for performing the biogeochemical analysis. Laura Espinosa-Asuar and Erika Aguirre-Planter provided technical and logistical assistance during the project.

