## AdditionalFile1 for "Population genomics of Vibrionaceae isolated from an endangered oasis reveals local adaptation after an environmental perturbation"

**Table of Contents:**

| **Supplementary Table 1. Nutrient measurements.** | 3 |
| --- | --- |
| **Supplementary Table 2. Nutrient concentrations ratio.** | 4 |
| **Supplementary Table 3. RDP classification of 16S rRNA.** | 5 |
| **Supplementary Table 4. Genomes general information.** | 13 |
| **Supplementary Table 5. Results of BUSCO analysis.** | 16 |
| **Supplementary Table 6. Results of recombination on coding genes.** | (Excel sheet) |
| **Supplementary Table 7. Metrics of clusters/gene families under selection.** | (Excel sheet) |
| **Supplementary Table 8. Genes found within genome regions found with GWAS analysis.** | (Excel sheet) |
| **Supplementary Table 9. GO term annotations of unique genes.** | 19 |
| **Supplementary Table 10. Illumina and 454 reads information and assembly processing.** | (Excel sheet) |
| **Supplementary Figure 1. Phylogenetic reconstruction of 16S rRNA.** | 20 |
| **Supplementary Figure 2. AdaptML ananalysis.** | 21 |
| **Supplementary Figure 3. Analysis of alpha values.** | 22 |
| **Supplementary Figure 4. Substructure of clade II.** | 23 |
| **Supplementary Figure 5. Frequency of recombination events.** | 24 |
| **Supplementary Figure 6. Recombination across genome.** | 25 |
| **Supplementary Figure 7. Membership of strains.** | 26 |

**Supplementary table 1**. Nutrient measures across sampled points

| Sediments | | | COT | NT | PT |
| --- | --- | --- | --- | --- | --- |
|  |  |  | mg g-1 | mg g-1 | mg g-1 |
| Ponds | Coordinates | |  |  |  |
| PR1 | N 26 52 16.7 | W 102 01 9.9 | 20.300 | 1.669 | 0.079 |
| PR2 | N 26 52 28.3 | W 102 01 18.8 | 35.600 | 3.966 | 0.197 |
| PR3 | N 26 52 8.2 | W 102 01 16.4 | 33.400 | 2.826 | 0.105 |
| PR4 | N 26 52 18.8 | W 102 01 16.7 | 23.300 | 1.723 | 0.093 |
| PR5 | N 26 52 18.1 | W 102 01 17.5 | 12.400 | 1.388 | 0.059 |
| PR6 | N 26 52 17.5 | W 102 01 18.6 | 45.200 | 5.112 | 0.225 |
| PR7 | N 26 52 17.7 | W 102 01 19.1 | 32.600 | 3.151 | 0.145 |
| PR8 | N 26 52 16.2 | W 102 01 18.6 | 30.000 | 3.434 | 0.247 |
| PR9 | N 26 52 15.7 | W 102 01 15.8 | 11.100 | 0.912 | 0.038 |

| Water | | | COT | NT | PT |
| --- | --- | --- | --- | --- | --- |
| Ponds | Coordinates | | mg l-1 | mg l-1 | mg l-1 |
| PR1 | N 26 52 16.7 | W 102 01 9.9 | 408.53 | 11.33 | 1.16 |
| PR2 | N 26 52 28.3 | W 102 01 18.8 | 448.73 | 16.16 | 1.38 |
| PR3 | N 26 52 8.2 | W 102 01 16.4 | 480.71 | 10.17 | 0.98 |
| PR4 | N 26 52 18.8 | W 102 01 16.7 | 186.23 | 5.08 | 1.15 |
| PR5 | N 26 52 18.1 | W 102 01 17.5 | 158.38 | 1.63 | 0.90 |
| PR6 | N 26 52 17.5 | W 102 01 18.6 | 499.55 | 11.70 | 1.27 |
| PR7 | N 26 52 17.7 | W 102 01 19.1 | 133.46 | 2.98 | 1.13 |
| PR8 | N 26 52 16.2 | W 102 01 18.6 | 247.09 | 7.42 | 1.10 |
| PR9 | N 26 52 15.7 | W 102 01 15.8 | 109.88 | 2.32 | 1.16 |

**Supplementary table 2.** Ratio of nutrients concentrations.

| Sediment | | | |
| --- | --- | --- | --- |
| Ponds | C:N | C:P | N:P |
| PR1 | 12.16 | 258.41 | 21.25 |
| PR2 | 8.98 | 180.81 | 20.14 |
| PR3 | 11.82 | 316.91 | 26.81 |
| PR4 | 13.53 | 249.71 | 18.46 |
| PR5 | 8.94 | 211.15 | 23.63 |
| PR6 | 8.84 | 200.54 | 22.68 |
| PR7 | 10.35 | 225.25 | 21.77 |
| PR8 | 8.74 | 121.35 | 13.89 |
| PR9 | 12.17 | 289.57 | 23.79 |

| Ponds | C:N | C:P | N:P |
| --- | --- | --- | --- |
| PR1 | 36.05101 | 350.9708 | 9.735395 |
| PR2 | 27.76795 | 324.6961 | 11.6932 |
| PR3 | 47.25816 | 489.5214 | 10.35845 |
| PR4 | 36.63061 | 161.658 | 4.413194 |
| PR5 | 97.04657 | 175.9778 | 1.813333 |
| PR6 | 42.69658 | 393.9669 | 9.227129 |
| PR7 | 44.84543 | 117.8975 | 2.628975 |
| PR8 | 33.30952 | 223.8134 | 6.719203 |
| PR9 | 47.28055 | 94.5611 | 2 |

**Supplementary table 3.** RDP classification of partial 16S rRNA. For each pond and environment, we obtained bacteria isolates, from which we got the partial sequence of the 16S rRNA gene (See methods for more information). Identification of each 16S rRNA sequence was performed with the classifier tool (Wang *et al.,* 2007).

| Isolate ID | Phylum | Class | Order | Family | Genus | Confidence |
| --- | --- | --- | --- | --- | --- | --- |
| PR2A33P | "Proteobacteria" | Gammaproteobacteria | "Enterobacteriales" | Enterobacteriaceae | Yersinia | 0.57 |
| PR4A1P | "Proteobacteria" | Gammaproteobacteria | "Enterobacteriales" | Enterobacteriaceae | Yersinia | 0.5 |
| PR6S15T | "Proteobacteria" | Gammaproteobacteria | "Enterobacteriales" | Enterobacteriaceae | Yersinia | 0.57 |
| PR9A22P | "Proteobacteria" | Gammaproteobacteria | "Enterobacteriales" | Enterobacteriaceae | Yersinia | 0.57 |
| PR1A41T | "Proteobacteria" | Gammaproteobacteria | "Vibrionales" | Vibrionaceae | Vibrio | 0.88 |
| PR1A60T | "Proteobacteria" | Gammaproteobacteria | "Vibrionales" | Vibrionaceae | Vibrio | 0.81 |
| PR1A64T | "Proteobacteria" | Gammaproteobacteria | "Vibrionales" | Vibrionaceae | Vibrio | 0.94 |
| PR1A73T | "Proteobacteria" | Gammaproteobacteria | "Vibrionales" | Vibrionaceae | Vibrio | 0.88 |
| PR1A8T | "Proteobacteria" | Gammaproteobacteria | "Vibrionales" | Vibrionaceae | Vibrio | 0.88 |
| PR1S1P | "Proteobacteria" | Gammaproteobacteria | "Vibrionales" | Vibrionaceae | Vibrio | 0.94 |
| PR1S1T | "Proteobacteria" | Gammaproteobacteria | "Vibrionales" | Vibrionaceae | Vibrio | 0.88 |
| PR1S2T | "Proteobacteria" | Gammaproteobacteria | "Vibrionales" | Vibrionaceae | Vibrio | 0.97 |
| PR1S30P | "Proteobacteria" | Gammaproteobacteria | "Vibrionales" | Vibrionaceae | Vibrio | 0.97 |
| PR1S3P | "Proteobacteria" | Gammaproteobacteria | "Vibrionales" | Vibrionaceae | Vibrio | 0.97 |
| PR1S3T | "Proteobacteria" | Gammaproteobacteria | "Vibrionales" | Vibrionaceae | Vibrio | 0.88 |
| PR1S4T | "Proteobacteria" | Gammaproteobacteria | "Vibrionales" | Vibrionaceae | Vibrio | 0.88 |
| PR1S50P | "Proteobacteria" | Gammaproteobacteria | "Vibrionales" | Vibrionaceae | Vibrio | 0.9 |
| PR1S5P | "Proteobacteria" | Gammaproteobacteria | "Vibrionales" | Vibrionaceae | Vibrio | 0.92 |
| PR1S5T | "Proteobacteria" | Gammaproteobacteria | "Vibrionales" | Vibrionaceae | Vibrio | 0.9 |
| PR1S6T | "Proteobacteria" | Gammaproteobacteria | "Vibrionales" | Vibrionaceae | Vibrio | 0.9 |
| PR2A1T | "Proteobacteria" | Gammaproteobacteria | "Vibrionales" | Vibrionaceae | Vibrio | 0.88 |
| PR2A25T | "Proteobacteria" | Gammaproteobacteria | "Vibrionales" | Vibrionaceae | Vibrio | 0.88 |
| PR2A27P | "Proteobacteria" | Gammaproteobacteria | "Vibrionales" | Vibrionaceae | Vibrio | 0.88 |
| PR2A34T | "Proteobacteria" | Gammaproteobacteria | "Vibrionales" | Vibrionaceae | Vibrio | 0.88 |
| PR2A8T | "Proteobacteria" | Gammaproteobacteria | "Vibrionales" | Vibrionaceae | Vibrio | 0.88 |
| PR2S10T | "Proteobacteria" | Gammaproteobacteria | "Vibrionales" | Vibrionaceae | Vibrio | 0.88 |
| PR2S12T | "Proteobacteria" | Gammaproteobacteria | "Vibrionales" | Vibrionaceae | Vibrio | 1 |
| PR2S17T | "Proteobacteria" | Gammaproteobacteria | "Vibrionales" | Vibrionaceae | Vibrio | 0.88 |
| PR2S21T | "Proteobacteria" | Gammaproteobacteria | "Vibrionales" | Vibrionaceae | Vibrio | 1 |
| PR2S30T | "Proteobacteria" | Gammaproteobacteria | "Vibrionales" | Vibrionaceae | Vibrio | 1 |
| PR2S4T | "Proteobacteria" | Gammaproteobacteria | "Vibrionales" | Vibrionaceae | Vibrio | 1 |
| PR3A10P | "Proteobacteria" | Gammaproteobacteria | "Vibrionales" | Vibrionaceae | Vibrio | 0.97 |
| PR3A14T | "Proteobacteria" | Gammaproteobacteria | "Vibrionales" | Vibrionaceae | Vibrio | 1 |
| PR3A28T | "Proteobacteria" | Gammaproteobacteria | "Vibrionales" | Vibrionaceae | Vibrio | 1 |
| PR3A8T | "Proteobacteria" | Gammaproteobacteria | "Vibrionales" | Vibrionaceae | Vibrio | 1 |
| PR3S12P | "Proteobacteria" | Gammaproteobacteria | "Vibrionales" | Vibrionaceae | Vibrio | 0.97 |
| PR3S12T | "Proteobacteria" | Gammaproteobacteria | "Vibrionales" | Vibrionaceae | Vibrio | 0.88 |
| PR3S1P | "Proteobacteria" | Gammaproteobacteria | "Vibrionales" | Vibrionaceae | Vibrio | 0.88 |
| PR3S1T1 | "Proteobacteria" | Gammaproteobacteria | "Vibrionales" | Vibrionaceae | Vibrio | 0.72 |
| PR3S1T | "Proteobacteria" | Gammaproteobacteria | "Vibrionales" | Vibrionaceae | Vibrio | 0.88 |
| PR3S34P | "Proteobacteria" | Gammaproteobacteria | "Vibrionales" | Vibrionaceae | Vibrio | 0.88 |
| PR3S3T | "Proteobacteria" | Gammaproteobacteria | "Vibrionales" | Vibrionaceae | Vibrio | 0.88 |
| PR3S5T | "Proteobacteria" | Gammaproteobacteria | "Vibrionales" | Vibrionaceae | Vibrio | 0.88 |
| PR3S9T | "Proteobacteria" | Gammaproteobacteria | "Vibrionales" | Vibrionaceae | Vibrio | 0.88 |
| PR4A10T | "Proteobacteria" | Gammaproteobacteria | "Vibrionales" | Vibrionaceae | Vibrio | 0.88 |
| PR4A19P | "Proteobacteria" | Gammaproteobacteria | "Vibrionales" | Vibrionaceae | Vibrio | 0.88 |
| PR4A1T | "Proteobacteria" | Gammaproteobacteria | "Vibrionales" | Vibrionaceae | Vibrio | 0.88 |
| PR4A23P | "Proteobacteria" | Gammaproteobacteria | "Vibrionales" | Vibrionaceae | Vibrio | 0.88 |
| PR4A5T | "Proteobacteria" | Gammaproteobacteria | "Vibrionales" | Vibrionaceae | Vibrio | 0.88 |
| PR4A6T | "Proteobacteria" | Gammaproteobacteria | "Vibrionales" | Vibrionaceae | Vibrio | 0.88 |
| PR4A8T | "Proteobacteria" | Gammaproteobacteria | "Vibrionales" | Vibrionaceae | Vibrio | 0.88 |
| PR4S10T | "Proteobacteria" | Gammaproteobacteria | "Vibrionales" | Vibrionaceae | Vibrio | 0.88 |
| PR4S1T | "Proteobacteria" | Gammaproteobacteria | "Vibrionales" | Vibrionaceae | Vibrio | 0.88 |
| PR4S3T | "Proteobacteria" | Gammaproteobacteria | "Vibrionales" | Vibrionaceae | Vibrio | 0.88 |
| PR4S5T | "Proteobacteria" | Gammaproteobacteria | "Vibrionales" | Vibrionaceae | Vibrio | 0.88 |
| PR4S6T | "Proteobacteria" | Gammaproteobacteria | "Vibrionales" | Vibrionaceae | Vibrio | 0.88 |
| PR4S8T | "Proteobacteria" | Gammaproteobacteria | "Vibrionales" | Vibrionaceae | Vibrio | 0.88 |
| PR2S5P | "Proteobacteria" | Gammaproteobacteria | "Vibrionales" | Vibrionaceae | Vibrio | 0.88 |
| PR3S6P | "Proteobacteria" | Gammaproteobacteria | "Vibrionales" | Vibrionaceae | Vibrio | 0.97 |
| PR7S5T | "Proteobacteria" | Gammaproteobacteria | "Vibrionales" | Vibrionaceae | Vibrio | 0.88 |
| PR5A7T | "Proteobacteria" | Gammaproteobacteria | "Vibrionales" | Vibrionaceae | Vibrio | 1 |
| PR5S13T | "Proteobacteria" | Gammaproteobacteria | "Vibrionales" | Vibrionaceae | Vibrio | 0.88 |
| PR5S5P | "Proteobacteria" | Gammaproteobacteria | "Vibrionales" | Vibrionaceae | Vibrio | 0.88 |
| PR6A28T1 | "Proteobacteria" | Gammaproteobacteria | "Vibrionales" | Vibrionaceae | Vibrio | 1 |
| PR6A28T2 | "Proteobacteria" | Gammaproteobacteria | "Vibrionales" | Vibrionaceae | Vibrio | 1 |
| PR6A3T1 | "Proteobacteria" | Gammaproteobacteria | "Vibrionales" | Vibrionaceae | Vibrio | 1 |
| PR6A6T2 | "Proteobacteria" | Gammaproteobacteria | "Vibrionales" | Vibrionaceae | Vibrio | 1 |
| PR6A8P1 | "Proteobacteria" | Gammaproteobacteria | "Vibrionales" | Vibrionaceae | Vibrio | 1 |
| PR6A8T | "Proteobacteria" | Gammaproteobacteria | "Vibrionales" | Vibrionaceae | Vibrio | 1 |
| PR6S12P | "Proteobacteria" | Gammaproteobacteria | "Vibrionales" | Vibrionaceae | Vibrio | 0.88 |
| PR6S14T2 | "Proteobacteria" | Gammaproteobacteria | "Vibrionales" | Vibrionaceae | Vibrio | 0.88 |
| PR6S15P | "Proteobacteria" | Gammaproteobacteria | "Vibrionales" | Vibrionaceae | Vibrio | 1 |
| PR6S20T | "Proteobacteria" | Gammaproteobacteria | "Vibrionales" | Vibrionaceae | Vibrio | 1 |
| PR6S34P | "Proteobacteria" | Gammaproteobacteria | "Vibrionales" | Vibrionaceae | Vibrio | 0.97 |
| PR6S7T | "Proteobacteria" | Gammaproteobacteria | "Vibrionales" | Vibrionaceae | Vibrio | 1 |
| PR7S17T | "Proteobacteria" | Gammaproteobacteria | "Vibrionales" | Vibrionaceae | Vibrio | 0.88 |
| PR7S7P | "Proteobacteria" | Gammaproteobacteria | "Vibrionales" | Vibrionaceae | Vibrio | 0.88 |
| PR8A3P | "Proteobacteria" | Gammaproteobacteria | "Vibrionales" | Vibrionaceae | Vibrio | 1 |
| PR8A6P | "Proteobacteria" | Gammaproteobacteria | "Vibrionales" | Vibrionaceae | Vibrio | 0.88 |
| PR8A6T | "Proteobacteria" | Gammaproteobacteria | "Vibrionales" | Vibrionaceae | Vibrio | 0.88 |
| PR8S9SAL | "Proteobacteria" | Gammaproteobacteria | "Vibrionales" | Vibrionaceae | Vibrio | 0.88 |
| PR9A10T | "Proteobacteria" | Gammaproteobacteria | "Vibrionales" | Vibrionaceae | Vibrio | 0.88 |
| PR9A1T | "Proteobacteria" | Gammaproteobacteria | "Vibrionales" | Vibrionaceae | Vibrio | 0.88 |
| PR9A5T | "Proteobacteria" | Gammaproteobacteria | "Vibrionales" | Vibrionaceae | Vibrio | 0.88 |
| PR9A6T | "Proteobacteria" | Gammaproteobacteria | "Vibrionales" | Vibrionaceae | Vibrio | 0.88 |
| PR9A9T | "Proteobacteria" | Gammaproteobacteria | "Vibrionales" | Vibrionaceae | Vibrio | 0.88 |
| PR9S7T | "Proteobacteria" | Gammaproteobacteria | "Vibrionales" | Vibrionaceae | Vibrio | 0.83 |
| PR3A3PO | "Proteobacteria" | Gammaproteobacteria | "Vibrionales" | Vibrionaceae | Vibrio | 1 |
| PR3A5PO | "Proteobacteria" | Gammaproteobacteria | "Vibrionales" | Vibrionaceae | Vibrio | 1 |
| PR4A1PO | "Proteobacteria" | Gammaproteobacteria | "Vibrionales" | Vibrionaceae | Vibrio | 1 |
| PR4A3PO | "Proteobacteria" | Gammaproteobacteria | "Vibrionales" | Vibrionaceae | Vibrio | 1 |
| PR4A7PO | "Proteobacteria" | Gammaproteobacteria | "Vibrionales" | Vibrionaceae | Vibrio | 1 |
| PR2S1PM | "Proteobacteria" | Gammaproteobacteria | "Vibrionales" | Vibrionaceae | Vibrio | 1 |
| PR2S13PM | "Proteobacteria" | Gammaproteobacteria | "Vibrionales" | Vibrionaceae | Vibrio | 1 |
| PR2S15PM | "Proteobacteria" | Gammaproteobacteria | "Vibrionales" | Vibrionaceae | Vibrio | 1 |
| PR2S17PM | "Proteobacteria" | Gammaproteobacteria | "Vibrionales" | Vibrionaceae | Vibrio | 1 |
| PR2S2PM | "Proteobacteria" | Gammaproteobacteria | "Vibrionales" | Vibrionaceae | Vibrio | 1 |
| PR2S24PM | "Proteobacteria" | Gammaproteobacteria | "Vibrionales" | Vibrionaceae | Vibrio | 1 |
| PR2S24PM2 | "Proteobacteria" | Gammaproteobacteria | "Vibrionales" | Vibrionaceae | Vibrio | 1 |
| PR2S5PM | "Proteobacteria" | Gammaproteobacteria | "Vibrionales" | Vibrionaceae | Vibrio | 1 |
| PR2S8PM | "Proteobacteria" | Gammaproteobacteria | "Vibrionales" | Vibrionaceae | Vibrio | 1 |
| PR1S10PM | "Proteobacteria" | Gammaproteobacteria | "Vibrionales" | Vibrionaceae | Vibrio | 1 |
| PR1S11PM | "Proteobacteria" | Gammaproteobacteria | "Vibrionales" | Vibrionaceae | Vibrio | 1 |
| PR1S14PM | "Proteobacteria" | Gammaproteobacteria | "Vibrionales" | Vibrionaceae | Vibrio | 1 |
| PR5S14PM | "Proteobacteria" | Gammaproteobacteria | "Vibrionales" | Vibrionaceae | Vibrio | 1 |
| PR7A18P | "Proteobacteria" | Gammaproteobacteria | Alteromonadales | Shewanellaceae | Shewanella | 1 |
| PR9A6P | "Proteobacteria" | Gammaproteobacteria | Alteromonadales | Shewanellaceae | Shewanella | 1 |
| PR3A8P | "Proteobacteria" | Gammaproteobacteria | "Enterobacteriales" | Enterobacteriaceae | Serratia | 0.94 |
| PR8A21T | "Proteobacteria" | Gammaproteobacteria | "Enterobacteriales" | Enterobacteriaceae | Serratia | 1 |
| PR8A5P | "Proteobacteria" | Gammaproteobacteria | "Enterobacteriales" | Enterobacteriaceae | Serratia | 1 |
| PR2A16PO | "Proteobacteria" | Gammaproteobacteria | "Enterobacteriales" | Enterobacteriaceae | Serratia | 1 |
| PR1S16PM | "Proteobacteria" | Gammaproteobacteria | "Enterobacteriales" | Enterobacteriaceae | Serratia | 0.94 |
| PR4S1PM | "Proteobacteria" | Gammaproteobacteria | "Enterobacteriales" | Enterobacteriaceae | Serratia | 1 |
| PR4S3PM1 | "Proteobacteria" | Gammaproteobacteria | "Enterobacteriales" | Enterobacteriaceae | Serratia | 1 |
| PR5A14PO | "Proteobacteria" | Gammaproteobacteria | Oceanospirillales | Halomonadaceae | Salinicola | 1 |
| PR5A15PO | "Proteobacteria" | Gammaproteobacteria | Oceanospirillales | Halomonadaceae | Salinicola | 1 |
| PR1A11P | "Proteobacteria" | Gammaproteobacteria | Chromatiales | Chromatiaceae | Rheinheimera | 1 |
| PR1A1P | "Proteobacteria" | Gammaproteobacteria | Chromatiales | Chromatiaceae | Rheinheimera | 1 |
| PR1S4P | "Proteobacteria" | Gammaproteobacteria | Chromatiales | Chromatiaceae | Rheinheimera | 1 |
| PR7S9P | "Proteobacteria" | Gammaproteobacteria | Alteromonadales | Pseudoalteromonadaceae | Pseudoalteromonas | 1 |
| PR2S1P | "Proteobacteria" | Gammaproteobacteria | "Vibrionales" | Vibrionaceae | Photobacterium | 1 |
| PR2S25P1 | "Proteobacteria" | Gammaproteobacteria | "Vibrionales" | Vibrionaceae | Photobacterium | 1 |
| PR2S44P | "Proteobacteria" | Gammaproteobacteria | "Vibrionales" | Vibrionaceae | Photobacterium | 1 |
| PR4S1P | "Proteobacteria" | Gammaproteobacteria | "Vibrionales" | Vibrionaceae | Photobacterium | 1 |
| PR4S2P | "Proteobacteria" | Gammaproteobacteria | "Vibrionales" | Vibrionaceae | Photobacterium | 1 |
| PR4S4P | "Proteobacteria" | Gammaproteobacteria | "Vibrionales" | Vibrionaceae | Photobacterium | 1 |
| PR1A2PO | "Proteobacteria" | Gammaproteobacteria | "Vibrionales" | Vibrionaceae | Photobacterium | 1 |
| PR3A19P | "Proteobacteria" | Gammaproteobacteria | Oceanospirillales | Oceanospirillaceae | Marinomonas | 1 |
| PR3A34P | "Proteobacteria" | Gammaproteobacteria | Oceanospirillales | Oceanospirillaceae | Marinomonas | 1 |
| PR1A31T | "Proteobacteria" | Gammaproteobacteria | "Vibrionales" | Vibrionaceae | Listonella | 0.67 |
| PR5S14T | "Proteobacteria" | Gammaproteobacteria | "Vibrionales" | Vibrionaceae | Listonella | 0.61 |
| PR1A18P | "Proteobacteria" | Gammaproteobacteria | Oceanospirillales | Halomonadaceae | Halomonas | 1 |
| PR8S6SAL | "Proteobacteria" | Gammaproteobacteria | Oceanospirillales | Halomonadaceae | Halomonas | 1 |
| PR8S8SAL | "Proteobacteria" | Gammaproteobacteria | Oceanospirillales | Halomonadaceae | Halomonas | 1 |
| PR4A11PO | "Proteobacteria" | Gammaproteobacteria | Oceanospirillales | Halomonadaceae | Halomonas | 1 |
| PR4A14PO | "Proteobacteria" | Gammaproteobacteria | Oceanospirillales | Halomonadaceae | Halomonas | 1 |
| PR4A15PO | "Proteobacteria" | Gammaproteobacteria | Oceanospirillales | Halomonadaceae | Halomonas | 1 |
| PR4A16PO | "Proteobacteria" | Gammaproteobacteria | Oceanospirillales | Halomonadaceae | Halomonas | 1 |
| PR4A17PO | "Proteobacteria" | Gammaproteobacteria | Oceanospirillales | Halomonadaceae | Halomonas | 1 |
| PR5A13PO | "Proteobacteria" | Gammaproteobacteria | Oceanospirillales | Halomonadaceae | Halomonas | 1 |
| PR5A16PO | "Proteobacteria" | Gammaproteobacteria | Oceanospirillales | Halomonadaceae | Halomonas | 1 |
| PR5A7PO | "Proteobacteria" | Gammaproteobacteria | Oceanospirillales | Halomonadaceae | Halomonas | 1 |
| PR6A1PO | "Proteobacteria" | Gammaproteobacteria | Oceanospirillales | Halomonadaceae | Halomonas | 1 |
| PR6A3PO | "Proteobacteria" | Gammaproteobacteria | Oceanospirillales | Halomonadaceae | Halomonas | 1 |
| PR5S11PM | "Proteobacteria" | Gammaproteobacteria | Oceanospirillales | Halomonadaceae | Halomonas | 1 |
| PR5S13PM | "Proteobacteria" | Gammaproteobacteria | Oceanospirillales | Halomonadaceae | Halomonas | 1 |
| PR1A4P | "Proteobacteria" | Gammaproteobacteria | "Enterobacteriales" | Enterobacteriaceae | Citrobacter | 0.97 |
| PR2A5T | "Proteobacteria" | Gammaproteobacteria | "Enterobacteriales" | Enterobacteriaceae | Citrobacter | 0.97 |
| PR1A15P | "Proteobacteria" | Gammaproteobacteria | Aeromonadales | Aeromonadaceae | Aeromonas | 1 |
| PR2A2P | "Proteobacteria" | Gammaproteobacteria | Aeromonadales | Aeromonadaceae | Aeromonas | 1 |
| PR2A41P | "Proteobacteria" | Gammaproteobacteria | Aeromonadales | Aeromonadaceae | Aeromonas | 1 |
| PR2S25P | "Proteobacteria" | Gammaproteobacteria | Aeromonadales | Aeromonadaceae | Aeromonas | 1 |
| PR2S7P | "Proteobacteria" | Gammaproteobacteria | Aeromonadales | Aeromonadaceae | Aeromonas | 1 |
| PR4A13P | "Proteobacteria" | Gammaproteobacteria | Aeromonadales | Aeromonadaceae | Aeromonas | 1 |
| PR4A7P | "Proteobacteria" | Gammaproteobacteria | Aeromonadales | Aeromonadaceae | Aeromonas | 1 |
| PR4S10P | "Proteobacteria" | Gammaproteobacteria | Aeromonadales | Aeromonadaceae | Aeromonas | 1 |
| PR4S8P | "Proteobacteria" | Gammaproteobacteria | Aeromonadales | Aeromonadaceae | Aeromonas | 1 |
| PR7A15P | "Proteobacteria" | Gammaproteobacteria | Aeromonadales | Aeromonadaceae | Aeromonas | 1 |
| PR7A16P | "Proteobacteria" | Gammaproteobacteria | Aeromonadales | Aeromonadaceae | Aeromonas | 1 |
| PR7A19P | "Proteobacteria" | Gammaproteobacteria | Aeromonadales | Aeromonadaceae | Aeromonas | 1 |
| PR7S1P | "Proteobacteria" | Gammaproteobacteria | Aeromonadales | Aeromonadaceae | Aeromonas | 1 |
| PR7S3P | "Proteobacteria" | Gammaproteobacteria | Aeromonadales | Aeromonadaceae | Aeromonas | 1 |
| PR7S4P | "Proteobacteria" | Gammaproteobacteria | Aeromonadales | Aeromonadaceae | Aeromonas | 1 |
| PR8A22P | "Proteobacteria" | Gammaproteobacteria | Aeromonadales | Aeromonadaceae | Aeromonas | 1 |
| PR8S5SAL | "Proteobacteria" | Gammaproteobacteria | Aeromonadales | Aeromonadaceae | Aeromonas | 1 |
| PR9A9P | "Proteobacteria" | Gammaproteobacteria | Aeromonadales | Aeromonadaceae | Aeromonas | 1 |
| PR9S11P | "Proteobacteria" | Gammaproteobacteria | Aeromonadales | Aeromonadaceae | Aeromonas | 1 |
| PR9S4P | "Proteobacteria" | Gammaproteobacteria | Aeromonadales | Aeromonadaceae | Aeromonas | 1 |
| PR6A5PO | "Proteobacteria" | Gammaproteobacteria | Aeromonadales | Aeromonadaceae | Aeromonas | 1 |
| PR6A9PO | "Proteobacteria" | Gammaproteobacteria | Aeromonadales | Aeromonadaceae | Aeromonas | 1 |
| PR8A13PO | "Proteobacteria" | Gammaproteobacteria | Aeromonadales | Aeromonadaceae | Aeromonas | 1 |
| PR8A14PO | "Proteobacteria" | Gammaproteobacteria | Aeromonadales | Aeromonadaceae | Aeromonas | 1 |
| PR7S2PM | "Proteobacteria" | Gammaproteobacteria | Aeromonadales | Aeromonadaceae | Aeromonas | 1 |
| PR3A1PO | "Proteobacteria" | Gammaproteobacteria | Pseudomonadales | Moraxellaceae | Acinetobacter | 1 |

**Supplementary table 4.** General information of the 42 Vibrionaceae genomes and 5 references used in this study.

| Clade | Strain | | Genome size (bp) | % GC | CDSs | BioProject/Seq Accession | BioSample | Isolation |
| --- | --- | --- | --- | --- | --- | --- | --- | --- |
| *Vibrio* II | *V. anguillarum* | 755 | 4,052,047 | 44.4 | 3,656 | PRJNA224116 | SAMN02603689 | Coho salmon (*Oncorhynchus kisutch*) |
| *Vibrio* II | *Vibrio sp.* | V01_P9A10T6 | 3,756,599 | 44.7 | 3,464 | PRJNA361510 | SAMN06234507 | CCB Los Hundidos |
| *Vibrio* II | *Vibrio sp.* | V02_P2A34T133 | 3,695,193 | 44.9 | 3,367 | PRJNA361510 | SAMN06251330 | CCB Los Hundidos |
| *Vibrio* II | *Vibrio sp.* | V03_P4A6T147 | 3,474,820 | 45.1 | 3,168 | PRJNA361511 | SAMN06251399 | CCB Los Hundidos |
| *Vibrio* II | *Vibrio sp.* | V04_P4A5T148 | 3,921,253 | 44.6 | 3,464 | PRJNA361511 | SAMN06251400 | CCB Los Hundidos |
| *Vibrio* II | *Vibrio sp.* | V05_P4A8T149 | 3,722,593 | 44.8 | 3,462 | PRJNA361511 | SAMN06251401 | CCB Los Hundidos |
| *Vibrio* II | *Vibrio sp.* | V06_P1A73T115 | 3,571,545 | 45 | 3,274 | PRJNA361511 | SAMN06251402 | CCB Los Hundidos |
| *Vibrio* II | *Vibrio sp.* | V07_P2A8T137 | 3,624,703 | 45 | 3,284 | PRJNA361511 | SAMN06251403 | CCB Los Hundidos |
| *Vibrio* II | *Vibrio sp.* | V08_P9A1T1 | 3,650,269 | 44.9 | 3,302 | PRJNA361511 | SAMN06251404 | CCB Los Hundidos |
| *Vibrio* II | *Vibrio sp.* | V09_P4A23P171 | 3,936,426 | 44.5 | 3,736 | PRJNA361511 | SAMN06251405 | CCB Los Hundidos |
| *Vibrio* II | *Vibrio sp.* | V10_P2A27P122 | 3,803,588 | 44.7 | 3,502 | PRJNA361511 | SAMN06251406 | CCB Los Hundidos |
| *Vibrio* II | *Vibrio sp.* | V11_P1A41T118 | 3,624,696 | 44.9 | 3,368 | PRJNA361511 | SAMN06251407 | CCB Los Hundidos |
| *Vibrio* II | *Vibrio sp.* | V12_P9A6T4 | 3,838,079 | 44.7 | 3,622 | PRJNA361511 | SAMN06251408 | CCB Los Hundidos |
| *Vibrio* II | *Vibrio sp.* | V14_P6S14T42 | 3,849,935 | 44.7 | 3,653 | PRJNA361511 | SAMN06251409 | CCB Los Hundidos |
| *Vibrio* II | *Vibrio sp.* | V15_P4S5T153 | 5,092,048 | 43.2 | 5,004 | PRJNA361511 | SAMN06251410 | CCB Los Hundidos |
| *Vibrio* II | *Vibrio sp.* | V17_P4S1T151 | 4,753,379 | 43.5 | 4,705 | PRJNA361511 | SAMN06251411 | CCB Los Hundidos |
| *Vibrio* II | *Vibrio sp.* | V18_P1S4T112 | 3,846,979 | 44.6 | 3,622 | PRJNA361511 | SAMN06251412 | CCB Los Hundidos |
| *Vibrio* II | *Vibrio sp.* | V19_P1S1T109 | 3,777,969 | 44.8 | 3,470 | PRJNA361511 | SAMN06251413 | CCB Los Hundidos |
| *Vibrio* II | *Vibrio sp.* | V20_P4S3T152 | 3,902,904 | 44.9 | 3,671 | PRJNA361511 | SAMN06251414 | CCB Los Hundidos |
| *Vibrio* II | *Vibrio sp.* | V22_P2S10T140 | 3,735,804 | 44.9 | 3,441 | PRJNA361511 | SAMN06251415 | CCB Los Hundidos |
| *Vibrio* II | *Vibrio sp.* | V23_P3S9T160 | 3,892,924 | 44.5 | 3,698 | PRJNA361511 | SAMN06251416 | CCB Los Hundidos |
| *Vibrio* II | *Vibrio sp.* | V24_P1S3T111 | 3,821,490 | 44.6 | 3,559 | PRJNA361511 | SAMN06251417 | CCB Los Hundidos |
| *Vibrio* II | *Vibrio sp.* | V25_P4S6T154 | 3,927,806 | 44.9 | 3,696 | PRJNA361511 | SAMN06251418 | CCB Los Hundidos |
| *Vibrio* III | *V. metschnikovii* | CIP 69.14 | 3,815,101 | 44.1 | 3,186 | PRJNA224116 | SAMN02393820 | Type strain |
| *Vibrio* III | *Vibrio sp.* | V31_P5A7T61 | 3,689,628 | 44.1 | 3,376 | PRJNA361511 | SAMN06251424 | CCB Los Hundidos |
| *Vibrio* III | *Vibrio sp.* | V32_P6A28T40 | 3,646,568 | 44.2 | 3,337 | PRJNA361511 | SAMN06251425 | CCB Los Hundidos |
| *Vibrio* III | *Vibrio sp.* | V33_P6A3T137 | 3,610,172 | 44.4 | 3,333 | PRJNA361511 | SAMN06251426 | CCB Los Hundidos |
| *Vibrio* III | *Vibrio sp.* | V34_P3A8T189 | 3,680,368 | 44.4 | 3,414 | PRJNA361511 | SAMN06251427 | CCB Los Hundidos |
| *Vibrio* III | *Vibrio sp.* | V40_P2S30T141 | 3,492,560 | 44.6 | 3,173 | PRJNA361511 | SAMN06251432 | CCB Los Hundidos |
| *Vibrio* IV | *Vibrio sp.* | V26_P1S5P106 | 3,121,437 | 43.9 | 2,887 | PRJNA361511 | SAMN06251419 | CCB Los Hundidos |
| *Vibrio* IV | *Vibrio sp.* | V27_P1S3P104 | 3,526,383 | 43.4 | 3,297 | PRJNA361511 | SAMN06251420 | CCB Los Hundidos |
| *Vibrio* IV | *Vibrio sp.* | V28_P6S34P95 | 3,503,965 | 43.5 | 3,244 | PRJNA361511 | SAMN06251421 | CCB Los Hundidos |
| *Vibrio* IV | *Vibrio sp.* | V29_P1S30P107 | 3,292,482 | 43.7 | 3,053 | PRJNA361511 | SAMN06251422 | CCB Los Hundidos |
| *Vibrio* IV | *Vibrio sp.* | V30_P3S12P165 | 3,524,720 | 43.4 | 3,266 | PRJNA361511 | SAMN06251423 | CCB Los Hundidos |
| Photobacterium | *Photobacterium sp.* | P44_P4S2P179 | 4,572,902 | 51.2 | 4,389 | PRJNA361511 | SAMN06251436 | CCB Los Hundidos |
| Photobacterium | *Photobacterium sp.* | P45_P2S44P127 | 4,546,445 | 51.3 | 4,368 | PRJNA361511 | SAMN06251437 | CCB Los Hundidos |
| Photobacterium | *Photobacterium sp.* | P46_P4S1P180 | 4,566,757 | 51.3 | 4,406 | PRJNA361511 | SAMN06251438 | CCB Los Hundidos |
| *Vibrio* V | *Vibrio sp.* | V36_P2S2PM302 | 4,900,770 | 50.1 | 4,673 | PRJNA361511 | SAMN06251428 | CCB Los Hundidos |
| *Vibrio* V | *Vibrio sp.* | V37_P2S8PM304 | 4,825,922 | 50.3 | 4,602 | PRJNA361511 | SAMN06251429 | CCB Los Hundidos |
| *Vibrio* V | *Vibrio sp.* | V38_P2S17PM301 | 4,945,276 | 50 | 4,746 | PRJNA361511 | SAMN06251430 | CCB Los Hundidos |
| *Vibrio* V | *Vibrio sp.* | V39_P1S14PM300 | 4,653,839 | 50.4 | 4,386 | PRJNA361511 | SAMN06251431 | CCB Los Hundidos |
| *Vibrio* VI | *V. parahaemolyticus* | BB220p | 5,103,524 | 45.3 | 4,442 | [PRJNA224116](https://www.ncbi.nlm.nih.gov/bioproject/PRJNA224116) | [SAMN02604302](https://www.ncbi.nlm.nih.gov/biosample/SAMN02604302) | Bangladesh environmental isolate |
| *Vibrio* VI | *V. alginolyticus* | NBRC 15630 | 5,146,637 | 44.6 | 4,457 | PRJNA224116 | [SAMN02603463](https://www.ncbi.nlm.nih.gov/biosample/SAMN02603463) | Spoiled horse mackerel |
| *Vibrio* VI | *Vibrio sp.* | V41_P2S12T139 | 3,613,116 | 45.6 | 3,320 | PRJNA361511 | SAMN06251433 | CCB Los Hundidos |
| *Vibrio* VI | *Vibrio sp.* | V42_P2S4T144 | 4,776,336 | 45.1 | 4,383 | PRJNA361511 | SAMN06251434 | CCB Los Hundidos |
| *Vibrio* VI | *Vibrio sp.* | V43_P6S15P86 | 4,846,002 | 45 | 4,468 | PRJNA361511 | SAMN06251435 | CCB Los Hundidos |
| *Vibrio* | *V. furnissii* | NCTC 11218 | 4,937,540 | 50.6 | 4,434 | [PRJNA224116](https://www.ncbi.nlm.nih.gov/bioproject/PRJNA224116) | [SAMN02603955](https://www.ncbi.nlm.nih.gov/biosample/SAMN02603955) | Estuarine environment |

**Supplementary table 5.** Results of the Benchmarking Universal Single-Copy Orthologs (BUSCO) analysis (see methods) using 452 BUSCOS of 721 species of the Gamma-proteobacteria database.

| **Clade** | **Strain** | | **BUSCO** | | | | |
| --- | --- | --- | --- | --- | --- | --- | --- |
|  |  |  | **Complete BUSCOs** | **Complete and single-copy BUSCOs** | **Complete and duplicated BUSCOs** | **Fragmented BUSCOs** | **Missing BUSCOs** |
| *Vibrio* II | *V. anguillarum* | 755 | 444 | 443 | 1 | 6 | 2 |
| *Vibrio* II | *Vibrio sp.* | V01_P9A10T6 | 444 | 444 | 0 | 3 | 5 |
| *Vibrio* II | *Vibrio sp.* | V02_P2A34T133 | 442 | 442 | 0 | 2 | 8 |
| *Vibrio* II | *Vibrio sp.* | V03_P4A6T147 | 430 | 429 | 1 | 5 | 17 |
| *Vibrio* II | *Vibrio sp.* | V04_P4A5T148 | 450 | 450 | 0 | 1 | 1 |
| *Vibrio* II | *Vibrio sp.* | V05_P4A8T149 | 435 | 435 | 0 | 6 | 11 |
| *Vibrio* II | *Vibrio sp.* | V06_P1A73T115 | 441 | 441 | 0 | 2 | 9 |
| *Vibrio* II | *Vibrio sp.* | V07_P2A8T137 | 441 | 441 | 0 | 3 | 8 |
| *Vibrio* II | *Vibrio sp.* | V08_P9A1T1 | 441 | 441 | 0 | 5 | 6 |
| *Vibrio* II | *Vibrio sp.* | V09_P4A23P171 | 451 | 451 | 0 | 1 | 0 |
| *Vibrio* II | *Vibrio sp.* | V10_P2A27P122 | 443 | 443 | 0 | 5 | 4 |
| *Vibrio* II | *Vibrio sp.* | V11_P1A41T118 | 439 | 438 | 1 | 4 | 9 |
| *Vibrio* II | *Vibrio sp.* | V12_P9A6T4 | 449 | 449 | 0 | 1 | 2 |
| *Vibrio* II | *Vibrio sp.* | V14_P6S14T42 | 451 | 451 | 0 | 1 | 0 |
| *Vibrio* II | *Vibrio sp.* | V15_P4S5T153 | 451 | 448 | 3 | 1 | 0 |
| *Vibrio* II | *Vibrio sp.* | V17_P4S1T151 | 442 | 440 | 2 | 4 | 6 |
| *Vibrio* II | *Vibrio sp.* | V18_P1S4T112 | 451 | 451 | 0 | 1 | 0 |
| *Vibrio* II | *Vibrio sp.* | V19_P1S1T109 | 435 | 435 | 0 | 7 | 10 |
| *Vibrio* II | *Vibrio sp.* | V20_P4S3T152 | 448 | 447 | 1 | 1 | 3 |
| *Vibrio* II | *Vibrio sp.* | V22_P2S10T140 | 445 | 445 | 0 | 2 | 5 |
| *Vibrio* II | *Vibrio sp.* | V23_P3S9T160 | 448 | 446 | 2 | 1 | 3 |
| *Vibrio* II | *Vibrio sp.* | V24_P1S3T111 | 450 | 450 | 0 | 1 | 1 |
| *Vibrio* II | *Vibrio sp.* | V25_P4S6T154 | 448 | 447 | 1 | 1 | 3 |
| *Vibrio* III | *V. metschnikovii* | CIP 69.14 | 421 | 421 | 0 | 19 | 12 |
| *Vibrio* III | *Vibrio sp.* | V31_P5A7T61 | 450 | 449 | 1 | 2 | 0 |
| *Vibrio* III | *Vibrio sp.* | V32_P6A28T40 | 450 | 449 | 1 | 2 | 0 |
| *Vibrio* III | *Vibrio sp.* | V33_P6A3T137 | 447 | 446 | 1 | 2 | 3 |
| *Vibrio* III | *Vibrio sp.* | V34_P3A8T189 | 449 | 448 | 1 | 2 | 1 |
| *Vibrio* III | *Vibrio sp.* | V40_P2S30T141 | 440 | 439 | 1 | 5 | 7 |
| *Vibrio* IV | *Vibrio sp.* | V26_P1S5P106 | 408 | 408 | 0 | 7 | 37 |
| *Vibrio* IV | *Vibrio sp.* | V27_P1S3P104 | 442 | 442 | 0 | 5 | 5 |
| *Vibrio* IV | *Vibrio sp.* | V28_P6S34P95 | 448 | 448 | 0 | 2 | 2 |
| *Vibrio* IV | *Vibrio sp.* | V29_P1S30P107 | 433 | 433 | 0 | 4 | 15 |
| *Vibrio* IV | *Vibrio sp.* | V30_P3S12P165 | 450 | 450 | 0 | 2 | 0 |
| Photobacterium | *Photobacterium sp.* | P44_P4S2P179 | 450 | 449 | 1 | 1 | 1 |
| Photobacterium | *Photobacterium sp.* | P45_P2S44P127 | 445 | 445 | 0 | 3 | 4 |
| Photobacterium | *Photobacterium sp.* | P46_P4S1P180 | 443 | 443 | 0 | 2 | 7 |
| *Vibrio* V | *Vibrio sp.* | V36_P2S2PM302 | 451 | 445 | 6 | 1 | 0 |
| *Vibrio* V | *Vibrio sp.* | V37_P2S8PM304 | 448 | 443 | 5 | 3 | 1 |
| *Vibrio* V | *Vibrio sp.* | V38_P2S17PM301 | 452 | 446 | 6 | 0 | 0 |
| *Vibrio* V | *Vibrio sp.* | V39_P1S14PM300 | 432 | 426 | 6 | 5 | 15 |
| *Vibrio* VI | *V. parahaemolyticus* | BB220p | 438 | 434 | 4 | 4 | 10 |
| *Vibrio* VI | *V. alginolyticus* | NBRC 15630 | 449 | 445 | 4 | 3 | 0 |
| *Vibrio* VI | *Vibrio sp.* | V41_P2S12T139 | 346 | 346 | 0 | 18 | 88 |
| *Vibrio* VI | *Vibrio sp.* | V42_P2S4T144 | 421 | 417 | 4 | 6 | 25 |
| *Vibrio* VI | *Vibrio sp.* | V43_P6S15P86 | 441 | 438 | 3 | 5 | 6 |
| *Vibrio* | *V. furnissii* | NCTC 11218 | 432 | 423 | 9 | 14 | 6 |

**Supplementary table 9.** GO terms enriched at the unique genes of each clade.

|  | GO.ID | Term | Annotated | Significant | Expected | Fisher test with Bonferroni correction |
| --- | --- | --- | --- | --- | --- | --- |
| Clade I | GO:0030153 | bacteriocin immunity | 4 | 3 | 0.04 | 0.0075576 |
| Clade II | GO:0016998 | cell wall macromolecule catabolic process... | 78 | 5 | 0.13 | 0.00038544 |
| Clade II | GO:0009253 | peptidoglycan catabolic process | 124 | 5 | 0.21 | 0.0038544 |
| Clade II | GO:0015891 | siderophore transport | 136 | 5 | 0.23 | 0.0061028 |
| Clade III | GO:0009103 | lipopolysaccharide biosynthetic process | 208 | 5 | 0.12 | 0.00022498 |
| Clade IV | GO:0006810 | transport | 16621 | 57 | 31.07 | 0.0013668 |
| Clade IV | GO:0002101 | tRNA wobble cytosine modification | 41 | 4 | 0.08 | 0.0017688 |
| Clade IV | GO:0051391 | tRNA acetylation | 41 | 4 | 0.08 | 0.0017688 |
| Clade IV | GO:0000160 | phosphorelay signal transduction system | 3099 | 20 | 5.79 | 0.002412 |
| Clade IV | GO:0032328 | alanine transport | 121 | 5 | 0.23 | 0.0057888 |
| Clade IV | GO:0007165 | signal transduction | 5325 | 34 | 9.95 | 0.038592 |
| Clade V | GO:0002101 | tRNA wobble cytosine modification | 41 | 4 | 0.04 | 0.0001608 |
| Clade V | GO:0051391 | tRNA acetylation | 41 | 4 | 0.04 | 0.0001608 |
| Clade V | GO:0055085 | transmembrane transport | 5369 | 20 | 5.5 | 0.002412 |
| Clade V | GO:0016310 | phosphorylation | 2230 | 12 | 2.28 | 0.0045024 |

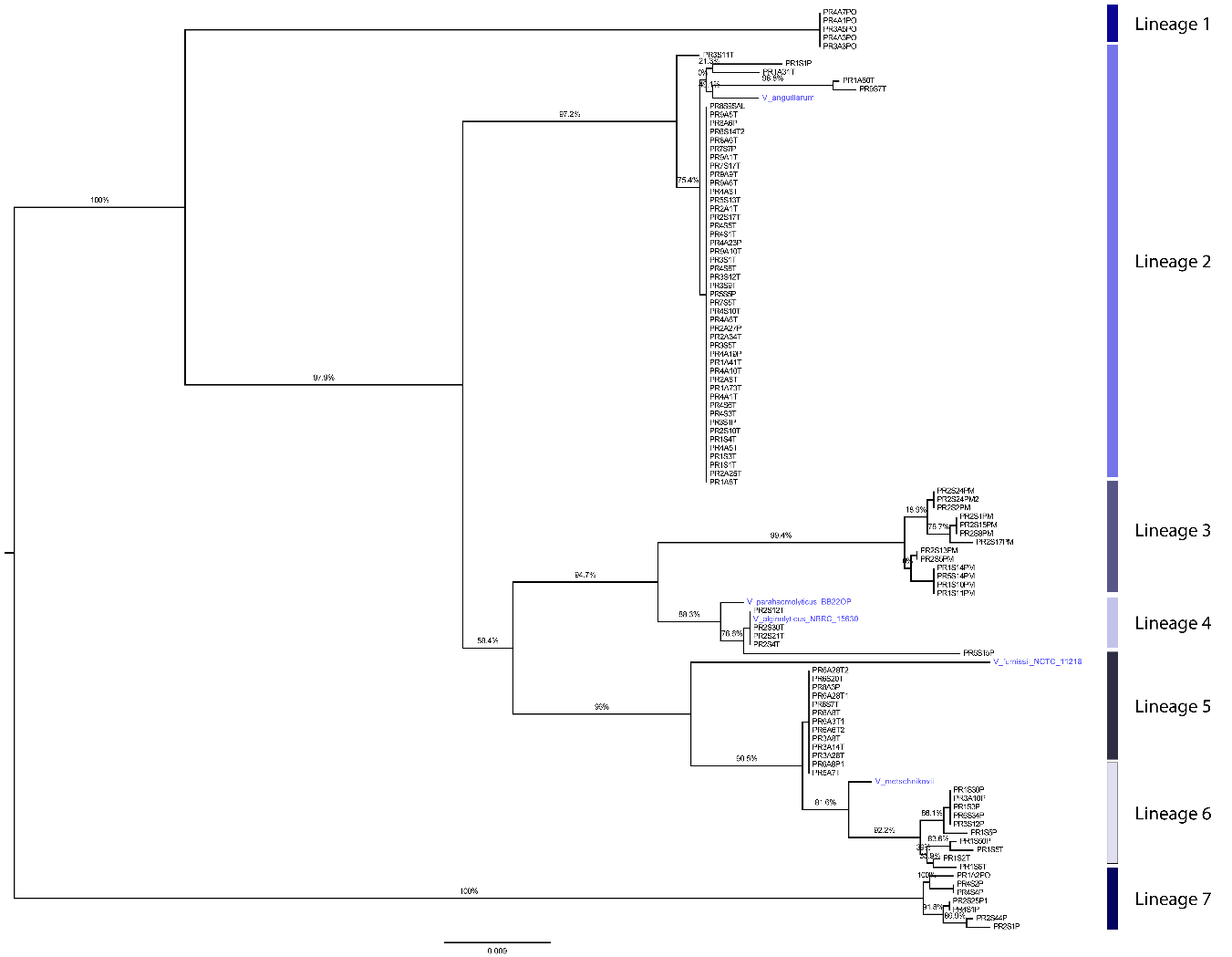

**Supplementary figure 1.** Phylogenetic reconstruction of 16S RNA sequences. A maximum likelihood method was employed over the first 700 bps of the 16S rRNA gene. Reference strains are shown in blue, as well seven lineages are highlighted with different colors. Strains for genome sequencing were selected from lineages, 2, 3, 4, 5, 6 and 7.

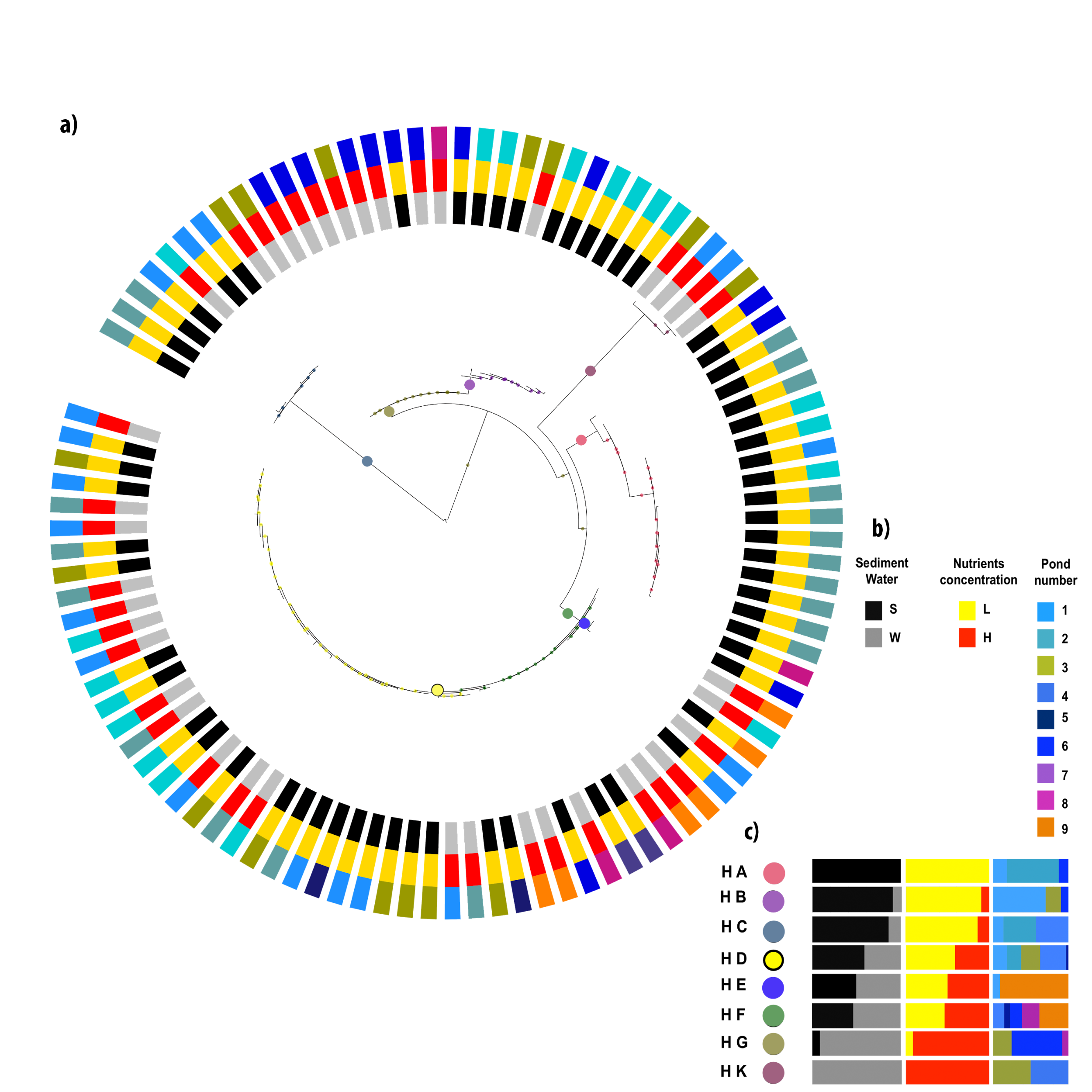

**Supplementary figure 2.** AdaptML analysis. Maximum likelihood phylogeny of partial 16S rRNA gene with *Halomonas* as outgroup. a) Shows the grouping of phylogroups given the environment from where they came from, colored dots within phylogeny indicates the projected habitat to which each phylogroup belong. b) Shows the color key for each environmental label. c) Shows the projected habitats and its specific content.

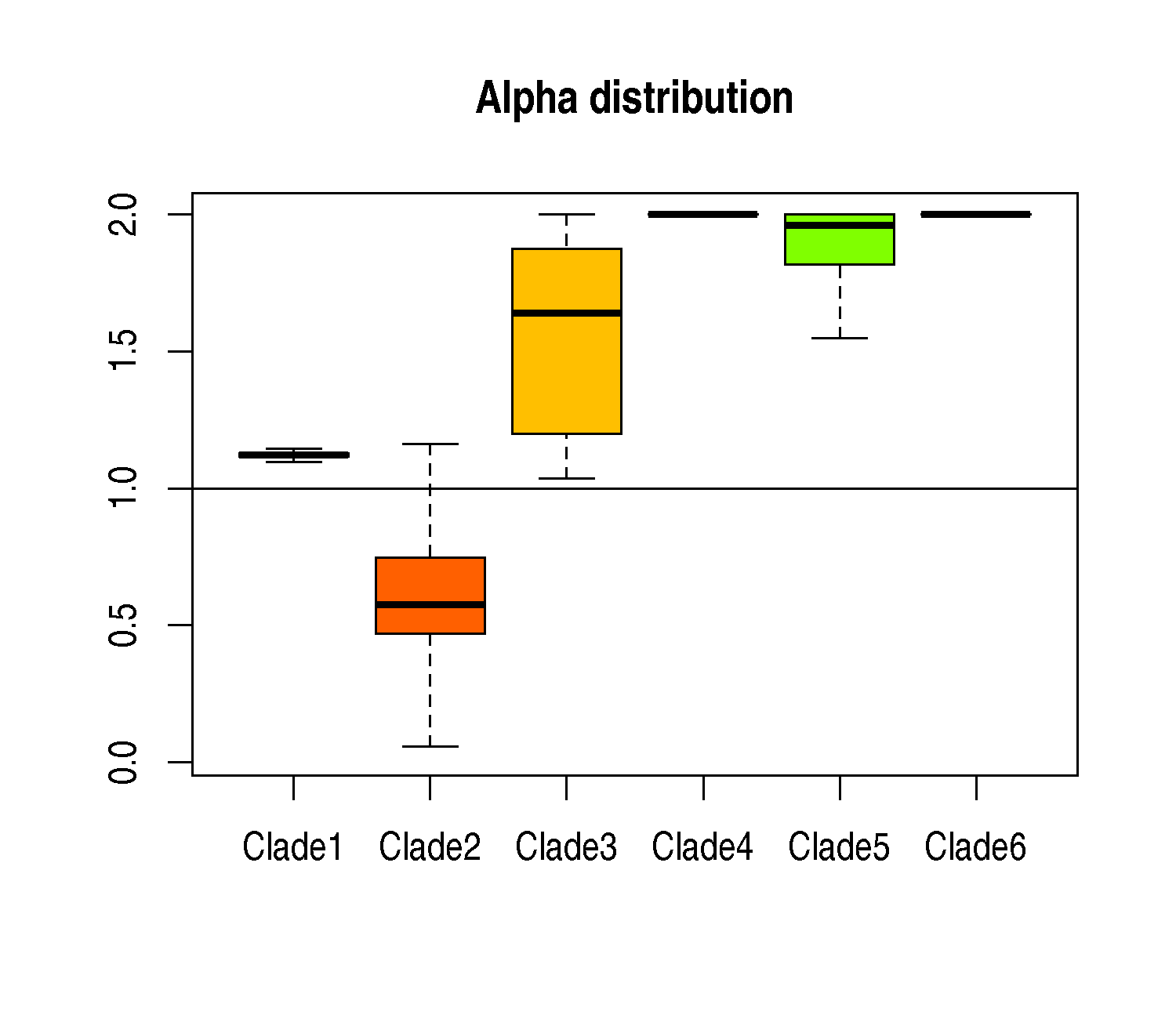

**Supplementary figure 3**. Random analysis of alpha values within each clade. The alpha values of each clade were calculated using three genomes, iterating this process 1000 types and changing randomly the three genomes each round of calculation. The “X” axis corresponds to each clade and “Y” axis to alpha values.

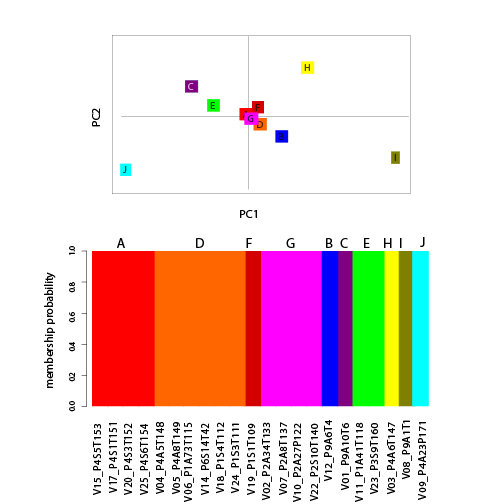

**Supplementary figure 4.** Analysis of structure of Clade II. The discriminant analysis of subgroups of Clade II displayed ten genetic groups (A-J), the strain composition of this genetic groups can be appreciated in the membership probability analysis below. The Sub-clades A, D and G were used for further analysis.

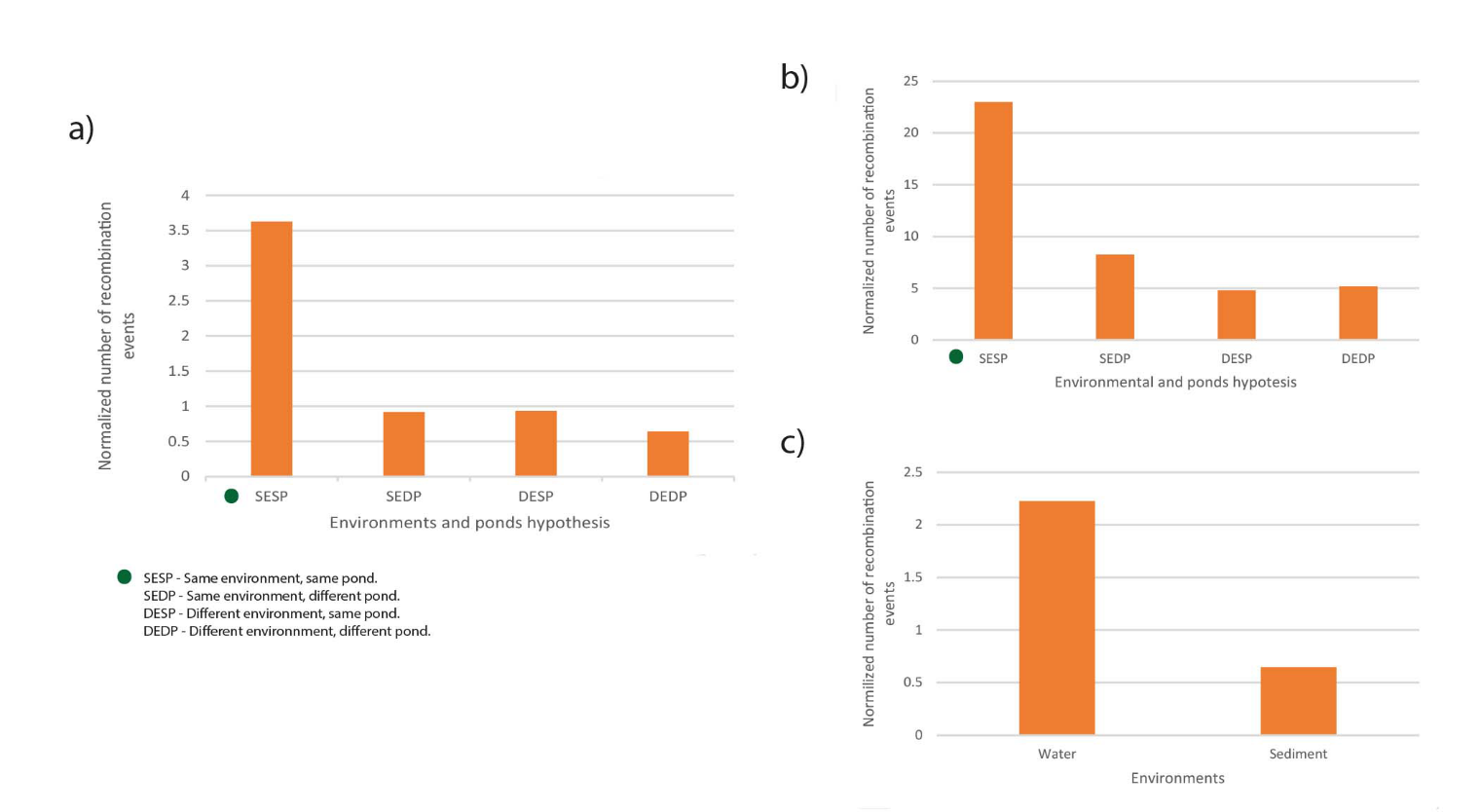

**Supplementary figure 5.** Frequency of recombination events. a) Shows frequency of recombination events of all Vibrionaceae. b) Shows frequency of recombination events based of Clade 2. c) Shows recombination events in terms of the origin of samples.

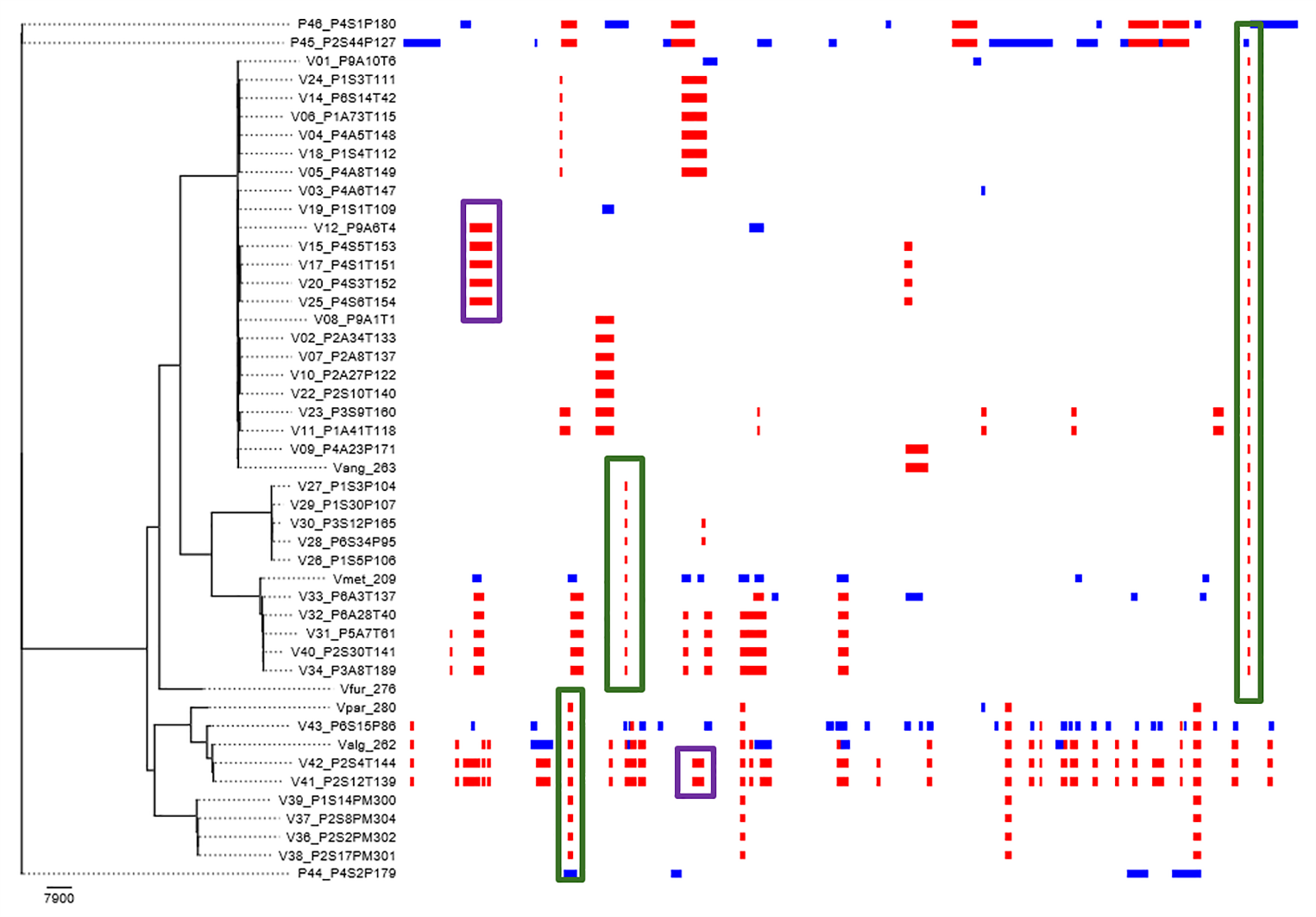

**Supplementary figure 6.** Recombination events across a whole genome alignment. Red blocks represent recombination events inside the alignment, while blue blocks represent points of recombination which origin was probably outside the alignment. Green boxed indicate sites that are shared with references and purple boxes indicate sites that had signals of recombination only within CCB strains.

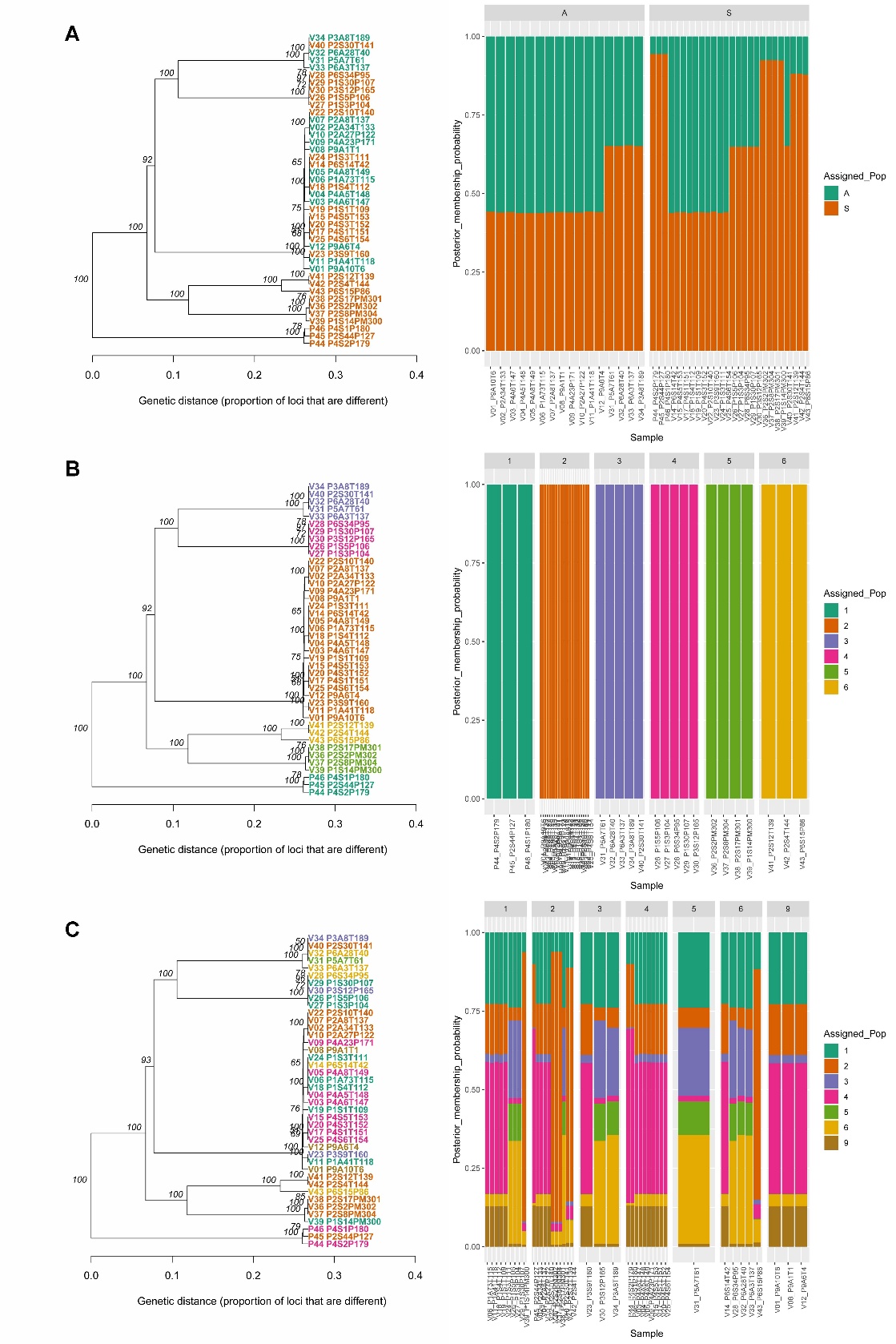

**Supplementary figure 7.** UPGMA and membership of the CCB strains. a) Clustering of the analyze strains according to their isolation environment. b) Clustering of the analyzed strains according to the predicted clade. c) Clustering of the analyzed strains according to their isolation pond.
